# A thiol–ene click-based strategy to customize injectable polymer–nanoparticle hydrogel properties for therapeutic delivery

**DOI:** 10.1101/2024.09.13.612978

**Authors:** Sophia J. Bailey, Noah Eckman, Elisa S. Brunel, Carolyn K. Jons, Samya Sen, Eric A. Appel

## Abstract

Polymer–nanoparticle (PNP) hydrogels are a promising injectable biomaterial platform that has been used for a wide range of biomedical applications including adhesion prevention, adoptive cell delivery, and controlled drug release. By tuning the chemical, mechanical, and erosion properties of injected hydrogel depots, additional control over cell compatibility and pharmaceutical release kinetics may be realized. Here, we employ thiol–ene click chemistry to prepare a library of modified hydroxypropylmethylcellulose (HPMC) derivatives for subsequent use in PNP hydrogel applications. When combined with poly(ethylene glycol)-b-poly(lactic acid) nanoparticles, we demonstrate that systematically altering the hydrophobic, steric, or pi stacking character of HPMC modifications can readily tailor the mechanical properties of PNP hydrogels. Additionally, we highlight the compatibility of the synthetic platform for the incorporation of cysteine-bearing peptides to access PNP hydrogels with improved bioactivity. Finally, through leveraging the tunable physical properties afforded by this method, we show hydrogel retention time *in vivo* can be dramatically altered without sacrificing mesh size or cargo diffusion rates. This work offers a route to optimize PNP hydrogels for a variety of translational applications and holds promise in the highly tunable delivery of pharmaceuticals and adoptive cells.

## 1. Introduction

Hydrogels are water-swollen soft materials with diverse biomedical applications ranging from regenerative medicine^1^ to drug delivery.^2^ Injectable hydrogel materials are particularly advantageous for minimally invasive delivery,^3^ where they have been employed for cell delivery^4,5^ and as reservoirs for the sustained delivery of pharmaceuticals.^6^ Several strategies to access injectable hydrogels exist, including *in situ* curing,^7^ microparticle hydrogel assemblies,^8^ dynamic covalent chemistries,^9,10^ and supramolecular interactions.^11^ Of these, polymer–nanoparticle (PNP) hydrogels are a promising class of supramolecular hydrogel that are formed by dynamic bridging interactions between polymers and nanoparticles. Our group has previously demonstrated that dodecyl-modified hydroxypropylmethylcellulose (HPMC-C12) can form supramolecular hydrophobic interactions with poly(ethylene glycol)-*b*-poly(lactic acid) nanoparticles (PEG-*b*-PLA NPs), providing injectable hydrogels well-suited for the sustained delivery of pharmaceuticals and cell-based therapies (**Figure 1a-b**).^12–14^ The physical properties of PNP hydrogels are a key consideration for designing materials for specific applications, as the stiffness, mesh size, and *in vivo* retention time of hydrogels are directly related to cell viability,^4,13^ cargo diffusion^2,15^ and pharmacokinetics,^2^ respectively.

**Figure 1.**
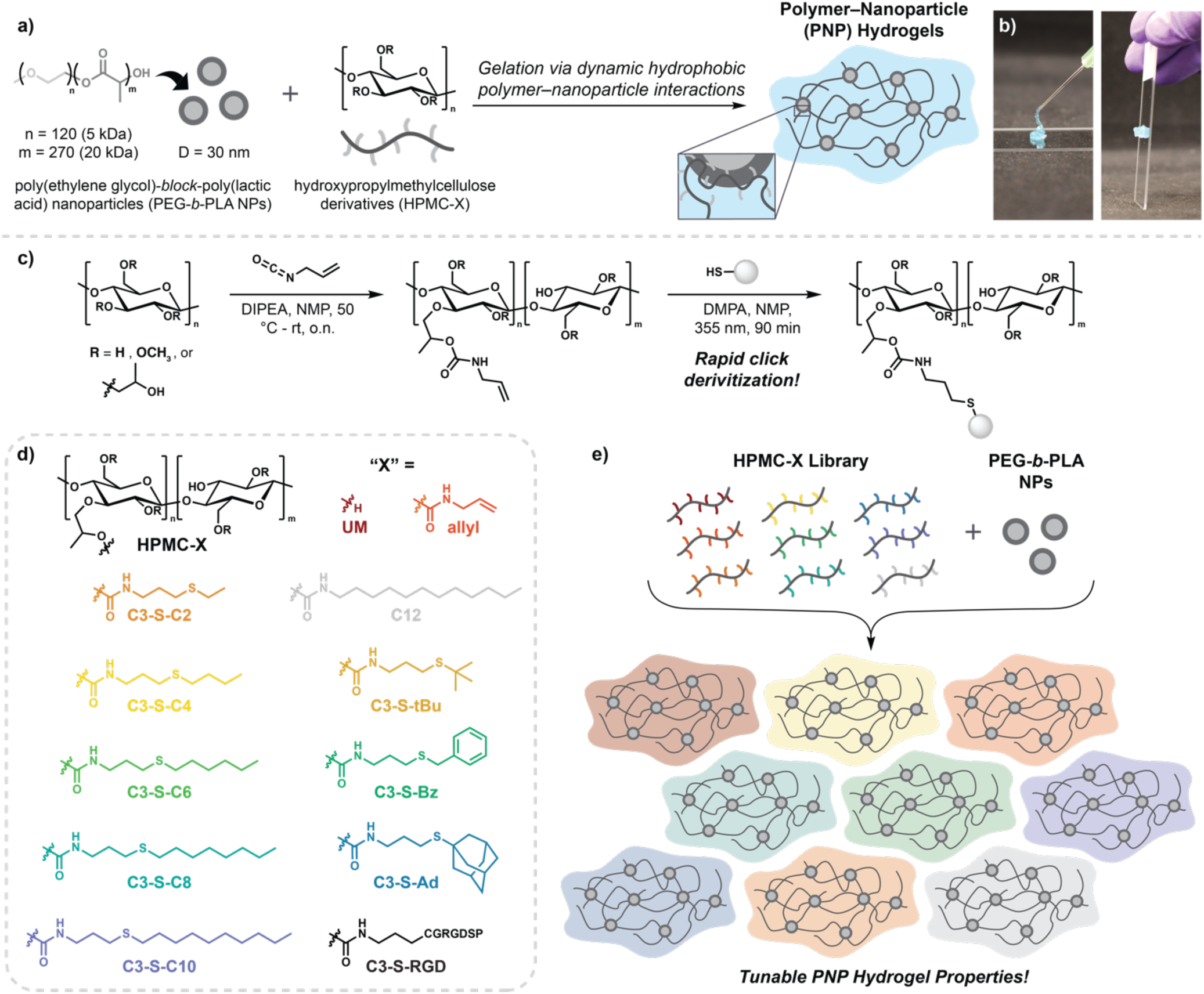
Preparation of PNP Hydrogel Library. a) General schematic of PNP hydrogel preparation. b) Photographs demonstrating injectability and recovery of PNP hydrogels. Blue color corresponds to a dye used for visualization only. c) Synthetic scheme to prepare HPMC-X derivatives. d) Library of HPMC-X derivatives prepared. e) Schematic depiction of PNP hydrogel library preparation.

To date, modulation of the physical properties of PNP hydrogels has largely been limited to altering the concentration of polymer and nanoparticles in solution. Thus, the stiffest, longest-lasting PNP hydrogels are accessed at high network densities, with the typical formulation being 2 wt% HPMC-C12 with 10 wt% PEG-PLA NPs, while softer, faster-dissolving hydrogels are prepared by lowering the concentrations of each component. Although much successful work has been demonstrated by this simple concentration-based method, this strategy fails to independently tailor hydrogel mechanics without changing network mesh size, and thus cargo diffusivity. Previous strategies to instead tune the strength of the supramolecular polymer–nanoparticle interaction have involved changing the identity of the nanoparticles, yielding unique hydrogel properties.^16^ However, this strategy is limited by the arduous synthetic requirements of block copolymer libraries and need for unique nanoprecipitation conditions. Thus, alternative chemical methods to tune PNP physical properties would enable better customization for broad biomedical applications.

We proposed that derivatization of HPMC might provide a more accessible route to tune polymer– nanoparticle interactions. Previous approaches to functionalize HPMC have relied upon reactions with isocyanates^12^ or electrostatic interactions,^17^ which are both limited by the commercial availability of potential pendant groups. Furthermore, isocyanates are highly reactive and suffer from poor shelf-stability, greatly limiting batch-to-batch reproducibility. Because of this, systematic studies comparing the effects of pendant group identity on polymer–nanoparticle interactions have been hindered by variable HPMC functionalization efficiencies. To thoroughly investigate pendant group effects on polymer–nanoparticle interactions and access therapeutic materials with new properties, an approach that provides consistent HPMC functionalization is needed. We envisioned click chemistry^18,19^ as a solution to this challenge, where a click handle could be installed on HPMC then easily derivatized via quantitative click reactions. With this strategy in mind, we selected thiol–ene click chemistry^20,21^ as an ideal candidate as it would allow for diverse derivatization with commercially available thiols and provide access to functionalization with cysteine-bearing peptides. Herein, we use this thiol–ene click strategy to investigate how systematically altering the hydrophobic, steric, or pi stacking character of HPMC modifications can be used to tailor the properties of subsequent PNP hydrogels. Notably, we demonstrate that PNP hydrogels can be prepared with consistent network density and diffusion properties while providing varied *in vivo* retention times, offering a new approach to access superior control over therapeutic cargo release.

## 2. Results and Discussion

### 2.1 Derivatization of HPMC

To access a uniformly functionalized series of HPMC derivatives, we opted to first prepare a large-batch parent material by treatment of HPMC with allyl isocyanate, followed by clicking to various thiol pendant groups (**Figure 1c**). For our library, we chose pendant groups which we hypothesized would readily impact the strength of the polymer–nanoparticle interaction, including thiols with varied alkyl chain length (C2 – C10); steric bulk (tBu and Ad) and competitive pi stacking interactions (Bz). To allow for comparison to previously reported HPMC-C12 PNP hydrogels, isocyanate feed ratio was maintained at 0.12 equivalents per cellulose repeat^12^ and incorporation of 5% relative to hydroxypropyl units was confirmed by ^1^H NMR (**Supporting Figure S1**). This single batch of parent HPMC-allyl was then exposed to various thiol–ene conditions to install our suite of pendant groups (**Figure 1d**). Successful HPMC functionalization was assessed by ^1^H NMR for each derivative, where consumption of allylic protons was confirmed and peaks corresponding to the various pendant functionalities could be readily observed (**Supporting Figure S2**).

Adjacently, this strategy offers a route to functionalize PNP hydrogels through the conjugation of cysteine-bearing peptides to HPMC. Previous routes to functionalize PNP hydrogels with peptides have relied on clicking peptides to PEG-*b*-PLA;^22^ however, these peptide-bearing block copolymers suffer from poor solubility, and can pose challenges in the subsequent nanoprecipitation. Thus, we envisioned the thiol– ene click strategy to provide an additional tool to introduce bioactivity to PNP hydrogels. To demonstrate the applicability of this alternative strategy for PNP functionalization, we conjugated a cell adhesive peptide, CRGDSP, to a 15% functionalized HPMC-allyl material, providing an RGD-dense biopolymer that could be introduced into PNP hydrogels. We found that doping this HPMC-RGD into PNP hydrogels for cell encapsulation led to improved cell morphology over traditional nonfunctionalized PNP hydrogels (**Supplemental Figure S4**). Although outside of the scope of this study, this functionalization strategy offers exciting potential to improve cell biocompatibility and elicit specific cellular behaviors through the incorporation of diverse peptides.

### 2.2 Rheological Properties of PNP Library

With a series of HPMC derivatives in hand, corresponding PNP hydrogels were prepared to investigate the effects of pendant group identity on rheological properties. Previous reports have demonstrated that increasing the hydrophobicity HPMC modifications greatly improves the strength of the supramolecular interactions with polymer nanoparticles, leading to stiffer materials.^12,23^ Based on these observations, we hypothesized that our library with well-matched functionalization would allow us to better detect nuances in the polymer–nanoparticle interactions via oscillatory shear rheometry.

Hydrogels were prepared by combining 1 wt% of each HPMC-X derivative with 5 wt% PEG-*b*-PLA NPs (**Figure 1e**) in phosphate buffered saline (PBS) (**Figure 1e**). For simplicity, PNP hydrogels are named after the “X” functionalization of the HPMC. Unmodified HPMC and dodecyl-modified HPMC following previously reported protocols^12^ were used for control PNP hydrogels denoted “UM” and “C12”, respectively. Upon mixing, all PNP derivatives form solid-like materials, with storage moduli (G’) greater than loss moduli (G”) across all frequencies tested (**Figure 2a-b, Supporting Figures S5–S15**). In contrast, without the addition of nanoparticles, frequency sweeps revealed liquid-like character for all 1 wt% solutions of HPMC-X derivatives (**Supporting Figures S5–S15**). We also prepared hydrogels out of nonfunctionalized HPMC exposed to the radical reaction conditions, and observed no changes in the rheological properties, highlighting the mild nature of the click conditions (**Supporting Figure S16**). Satisfyingly, we found that increasing pendant chain length led to a gradual increase in PNP hydrogel modulus likely due to increased interactions between the hydrophobic pendant groups and the PEG-*b*-PLA NPs. We also found that nuances between sterically bulky adamantyl group were easily detected in comparison to the linear C3-S-C10; although, no impact was noted when comparing the smaller C3-S-tBu and C3-S-C4 groups. A decrease in stiffness was also observed for C3-S-Bz groups over linear C3-S-C6, indicating competitive pi stacking between HPMC chains might reduce the overall number of polymer– nanoparticle interactions contributing to the material properties. In comparison to polymer–nanoparticle interactions, however, we found that pi stacking interactions alone did not appear to enhance the storage modulus (**Supporting Figure S17**). Interestingly, in amplitude sweeps of all PNP hydrogel derivatives, we observed characteristic G” peaks at the crossover with G’, indicative of a jamming transition between polymer–nanoparticle units.^24^ Observation of this transition had been reported for PNP derivatives prepared with HPMC-C12, but previous HPMC derivatization routes and rheological methods had precluded the observation of this phenomenon (**Supporting Figure S5–S15**).^23^

**Figure 2.**
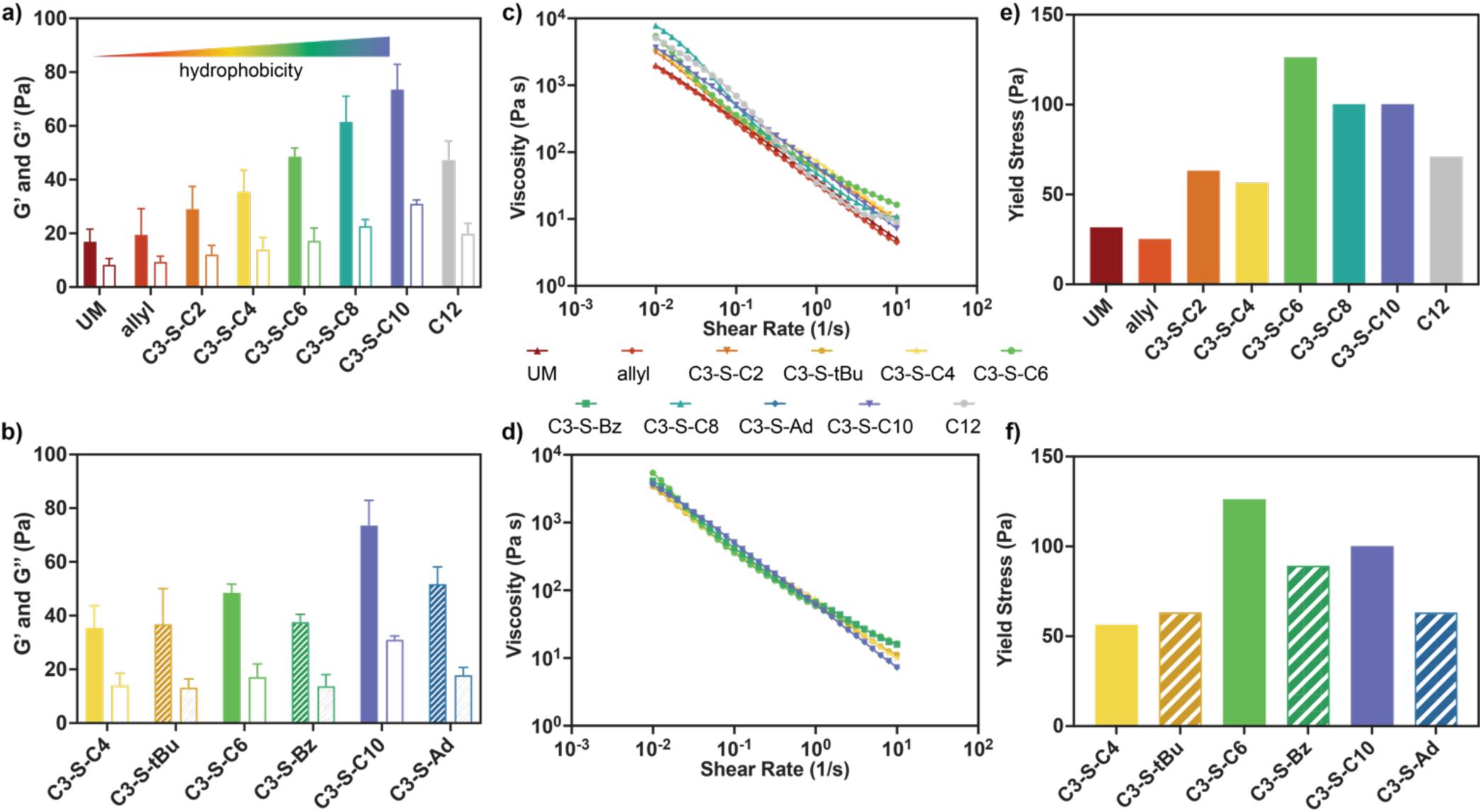
Rheological Properties of PNP Library. Average storage (G’, filled bars) and loss (G”, empty bars) moduli of a) linear and b) steric/pi stacking PNP derivatives at a representative angular frequency in the linear viscoelastic regime (ω = 1 rad/s, 2% strain). Values reported are means ± SE of n = 3. Steady shear rheology of c) linear and d) steric/pi stacking PNP derivatives. Yield stresses of e) linear and f) steric/pi stacking PNP derivatives determined from stress ramps where viscosity dropped below half of the initial plateau viscosity.

The ability for PNP hydrogels to be readily injected then form robust depots *in vivo* has enabled versatile biomedical applications to be realized; thus, we sought to confirm that these essential properties were preserved across the new PNP library. First, we conducted steady shear rheometry, where we were pleased to observe the viscosities for all materials decreased multiple orders of magnitude with increasing shear rates (**Figure 2c-d**). We then ran stress ramp measurements to determine the yield stress for each derivative, a predictor of depot persistence *in vivo*.^14,25^ Similar to the trends observed in moduli, we found that increased pendant hydrophobicity led to higher zero shear viscosities and yield stresses (**Figure 2e**, **Supporting Figure S19**), indicative of stronger interactions between nanoparticles with pendant groups of increasing chain lengths. Likewise, steric and pi stacking interference led to decreased zero shear viscosities and yield stresses compared to linear analogs (**Figure 2f**, **Supporting Figure S19**).

Flow sweeps and stress ramp measurements provided confirmation of the hydrogels’ ability to flow under high shear; however, we wished to further investigate the dynamics of the polymer–nanoparticle interactions which would be applicable to depots post-injection. Even after the cessation of high shear, dynamic supramolecular interactions allow PNP hydrogels to exhibit viscoelastic behaviors, which play important roles on depot persistence and cargo delivery when exposed to the *in vivo* environment.^25^ Given the observed differences in stiffness and shear-thinning properties across our PNP library, we predicted our stronger interactions would slow the rate of network dissociation and lead to observed differences in viscoelastic behaviors like stress relaxation.^26,27^ Stress relaxation experiments revealed that all PNP hydrogels readily relax stress (**Figure 3a-b**), with characteristic relaxation times on the order of 10^2^ – 10^3^ seconds that increased with pendant hydrophobicity (**Figure 3c-d**). Interestingly, C3-S-Bz pendants led to an unexpected increase in relaxation time, which could be attributed to breaking of pi stacking interactions. Fitting of frequency sweep data to continuous viscoelastic relaxation spectra was also conducted determine G’/G’’ crossovers for each PNP hydrogel derivative, revealing similar, but less dramatic trends compared to stress relaxation measurements (**Supporting Figures S21–22**). We next conducted creep tests to investigate material deformation under subcutaneously relevant stress of 20 Pa, an ideal predictor of depot flattening and retention for *in vivo* mouse models.^25^ We observed rapid deformation of PNP hydrogels prepared with unmodified HPMC and allyl pendants, while increasing pendant chain lengths again provided a satisfying trend, with the most hydrophobic C3-S-C10 material exhibiting the lowest compliance (**Figure 3e**). For sterically hindered and pi stacking pendants, notable differences in creep behavior were only observed between C3-S-Ad and its linear analog C3-S-C10 (**Figure 3f**). To test stress values relevant to other organisms, we also investigated the creep response of each PNP hydrogel derivative under applied stresses up to 100 or 200 Pa and found similar trends (**Supporting Figures S23–24**).

**Figure 3.**
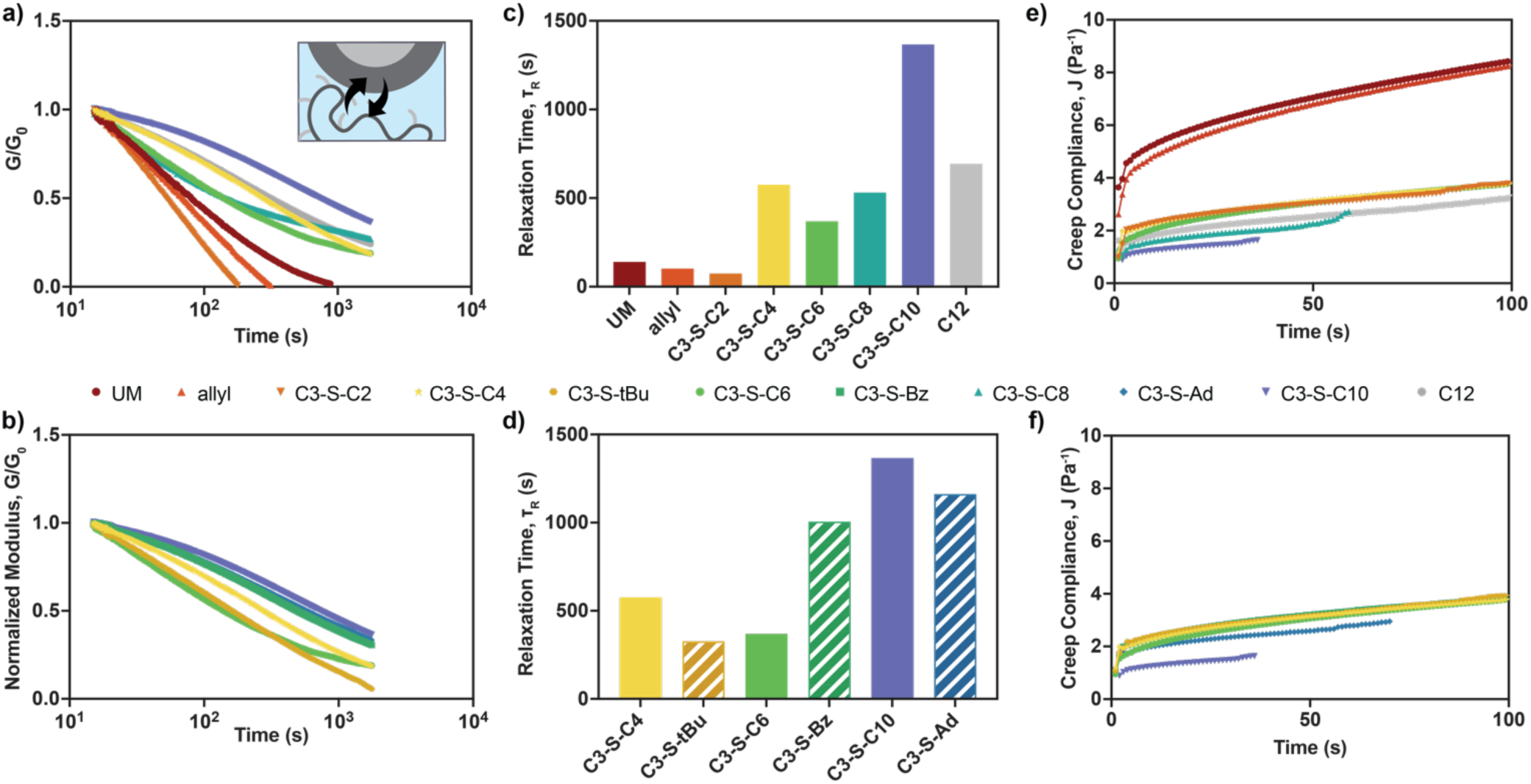
Dynamic Properties of PNP Library. Stress relaxation measurements of a) linear and b) steric/pi stacking PNP derivatives (5% strain). Relaxation times, τ_R_, of c) linear and d) steric/pi stacking PNP derivatives determined from fitting of relaxation curves in a) and b). Creep compliance of a) linear and b) steric/pi stacking PNP derivatives under a constant applied stress of 20 Pa.

In addition to probing network dissociation dynamics, we were also interested in the thixotropic behaviors of hydrogels: an important consideration for rapid depot formation after injection and prevention of burst release.^28^ To investigate these behaviors, we conducted stress overshoot experiments to obtain characteristic material recovery times. Stress overshoot experiments were conducted as previously described,^16^ where stress overshoots during shear were measured following increasing wait times (**Figure 4a-c**). Fitting of stress overshoots to an exponential plateau provides characteristic timescales for this healing behavior, τ_H_.^16,29^ These experiments revealed that long, linear pendant groups C3-S-C8, C3-S-C10 and C12 required nearly twice the time to heal as all other derivatives (**Figure 4d**). Small differences were also noted when comparing linear derivatives to their sterically hindered and pi stacking counterparts (**Figure 4e**). These results indicate that although these pendant groups ultimately form stronger polymer– nanoparticle interactions, the rate of association is also significantly slowed. This observation indicates that there could be an important tradeoff between material retention time and initial burst release due to slowed network recovery post-injection.

**Figure 4.**
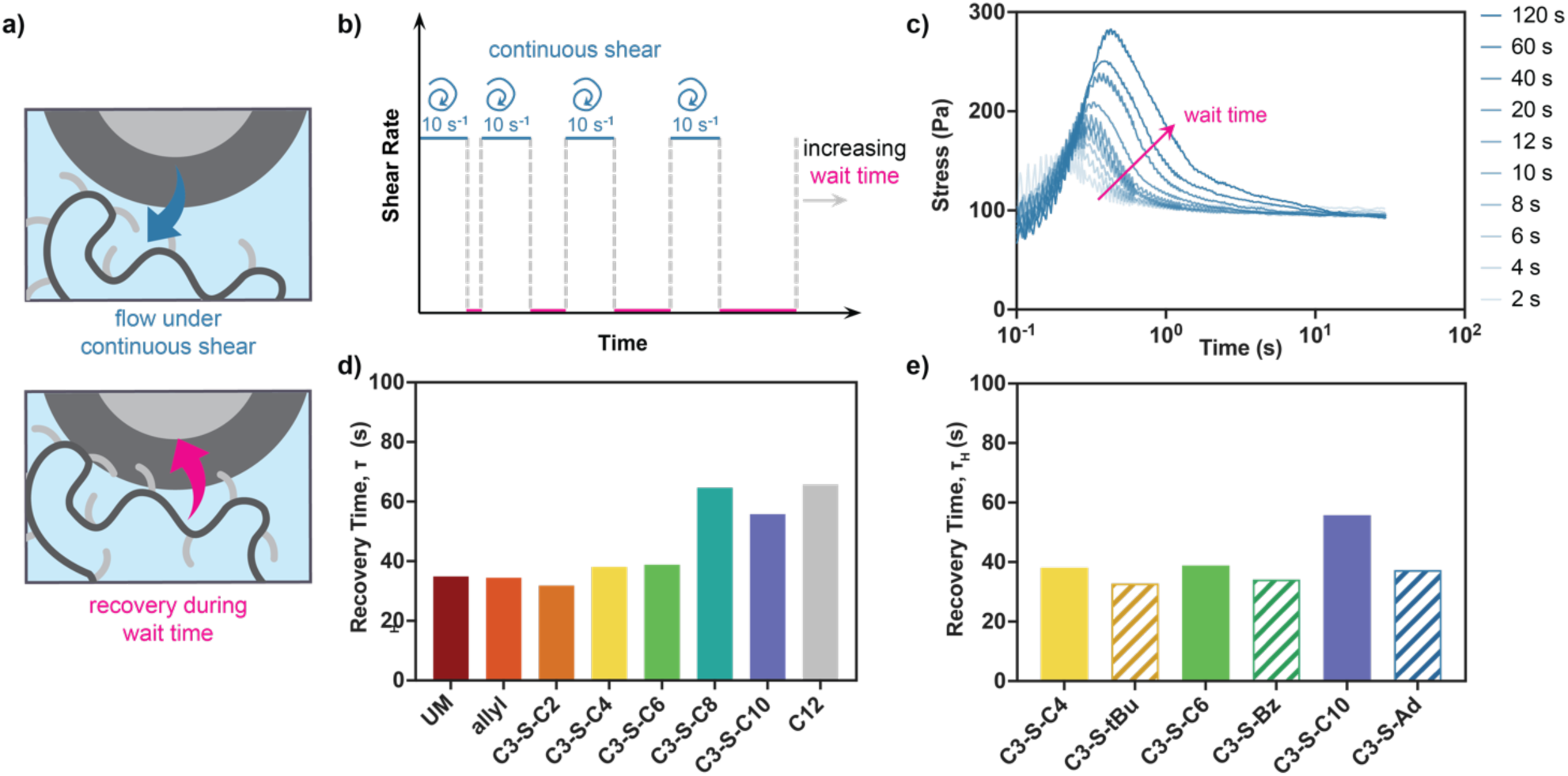
Recovery Properties of PNP Library. a) Simplified depiction of polymer–nanoparticle interaction under flow and wait time conditions. b) Diagram describing experimental protocol to determine recovery times. c) Representative stress overshoot measurement of an Ad PNP hydrogel with increasing wait times. Recovery times, τ_H_, of d) linear and e) steric/pi stacking PNP derivatives determined by fitting stress overshoots.

### 2.3 Cargo Diffusion and Release

Previous work has established that timescales of biotherapeutic cargo delivery can be readily tuned through altering PNP hydrogel network density;^15^ however, this strategy simultaneously changes both cargo diffusivity and hydrogel lifetime, limiting the overall control over cargo release. In contrast, we hypothesized that through chemical modification of the polymer–nanoparticle interaction we would effectively maintain mesh size and cargo diffusivity, providing control over exposure exclusively through material retention. To test this, we investigated the diffusivity of a model cargo, bovine albumin serum (BSA) across the PNP library (**Figure 5a**) using fluorescence recovery after photobleaching (FRAP) experiments. PNP hydrogels loaded with 0.05wt% fluorescein-labeled BSA as a model were prepared, and fluorescence recovery was monitored after bleaching with high intensity light. Diffusivity values for BSA within each PNP hydrogel derivative were then determined from these recovery curves. We found that across all derivatives, diffusion is more than an order of magnitude slower than calculated for PBS – indicative of successful entrapment within the hydrogel mesh (**Figure 5b**). Additionally, minimal differences in diffusivity were observed between the most mechanically distinct PNP hydrogels UM and C3-S-C10, indicating cargo release could be tuned selectively through material erosion properties rather than varied cargo diffusivity. To further investigate release, PNP hydrogels containing BSA conjugated to AlexaFluor 647 (0.025 wt%) were incubated in phosphate buffered saline at 37 °C for two weeks while media was monitored via fluorescence. As illustrated in **Figure 5c**, all derivatives possess similar release profiles with less than 30% release over two weeks. Together, these release experiments suggest that using this chemical modification strategy will allow for exquisite control over biotherapeutic cargo exposure purely through polymer–nanoparticle interaction strength, rather than network density.

**Figure 5.**
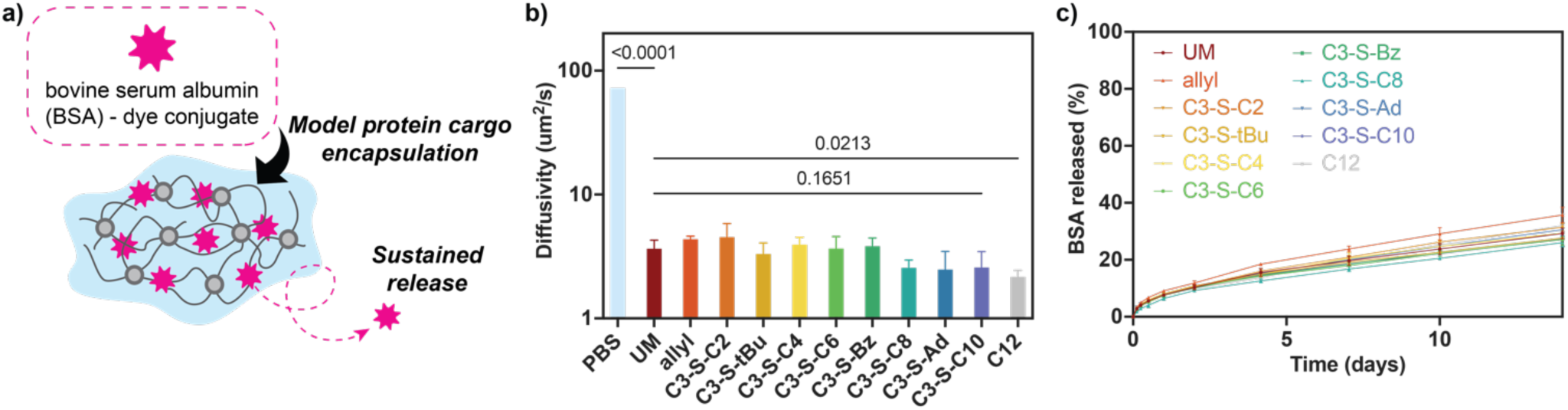
a) Schematic Depiction of Sustained Cargo Release. b) Bar chart plotting diffusivities of BSA-FITC determined by FRAP (n=3) for each PNP hydrogel derivative. PBS value calculated by Stokes-Einstein equation. Statistical analysis between groups was run using GraphPad Prism using an unpaired t-test with two-tails. c) *In vitro* release of BSA-AlexaFluor 647 from each PNP hydrogel derivative into PBS over two weeks at 37 °C. Values reported are means ± SE of n = 3.

### 2.4 *In Vivo* Material Retention

Having demonstrated that PNP hydrogel mechanical properties are directly related to the polymer– nanoparticle affinity through HPMC derivatization we proposed that we could also tune material retention *in vivo*. Four derivatives were chosen to test that spanned the range of physical properties obtained from the library: UM, C3-S-C4, C3-S-C10, and the previously reported C12 (**Figure 6a-c**). Female C57BL/6 mice were injected subcutaneously with PNP hydrogels and depots were measured with calipers at designated time points for 70 days, at which point mice were euthanized and remaining depots were excised to assess fibrosis and mass retention (**Figure 6d**). As highlighted in **Figure 6e-g**, creep and yielding behaviors remain a good predictor of *in vivo* retention.^25^ Notably, our UM hydrogels which exhibit the greatest creep compliance under subcutaneously relevant stresses led to the most rapid depot dissolution, with all five undetectable by day 46. The next most compliant material, C3-S-C4, had improved material retention, with only one depot still observable by the end of the 10-week study. In contrast, the most robust C3-S-C10 and C12 materials maintain observable depots through the endpoint of the study. At day 70, excisions of all remaining depots were conducted and revealed no fibrotic capsule formation (**Supporting Figure S27**). C3-S-C10 and C12 explants were easily removed with 9.2% and 11.4% mass retention, respectively (**Figure 6h**). In contrast, C3-S-C4 had lower mass retention of 4.1%, with only 3 detectable gels excised. Likewise, only one explant was obtained from the UM group, with an overall mass retention of 0.2%. This *in vivo* work highlights the tunability of PNP hydrogel retention times afforded by this platform, including access to depots that dissolve much faster than the previously reported HPMC-C12-based formulation,^12^ yet maintain low diffusivity as demonstrated in **Section 2.3**. Thus, this newfound control has the potential to provide well-matched dissolution rates with desired drug release profiles. Notably, faster dissolving hydrogels could also circumvent the need for multiple injection sites necessitated by hydrogel persistence beyond drug dosing regimens, a common challenge with depot technologies.

**Figure 6.**
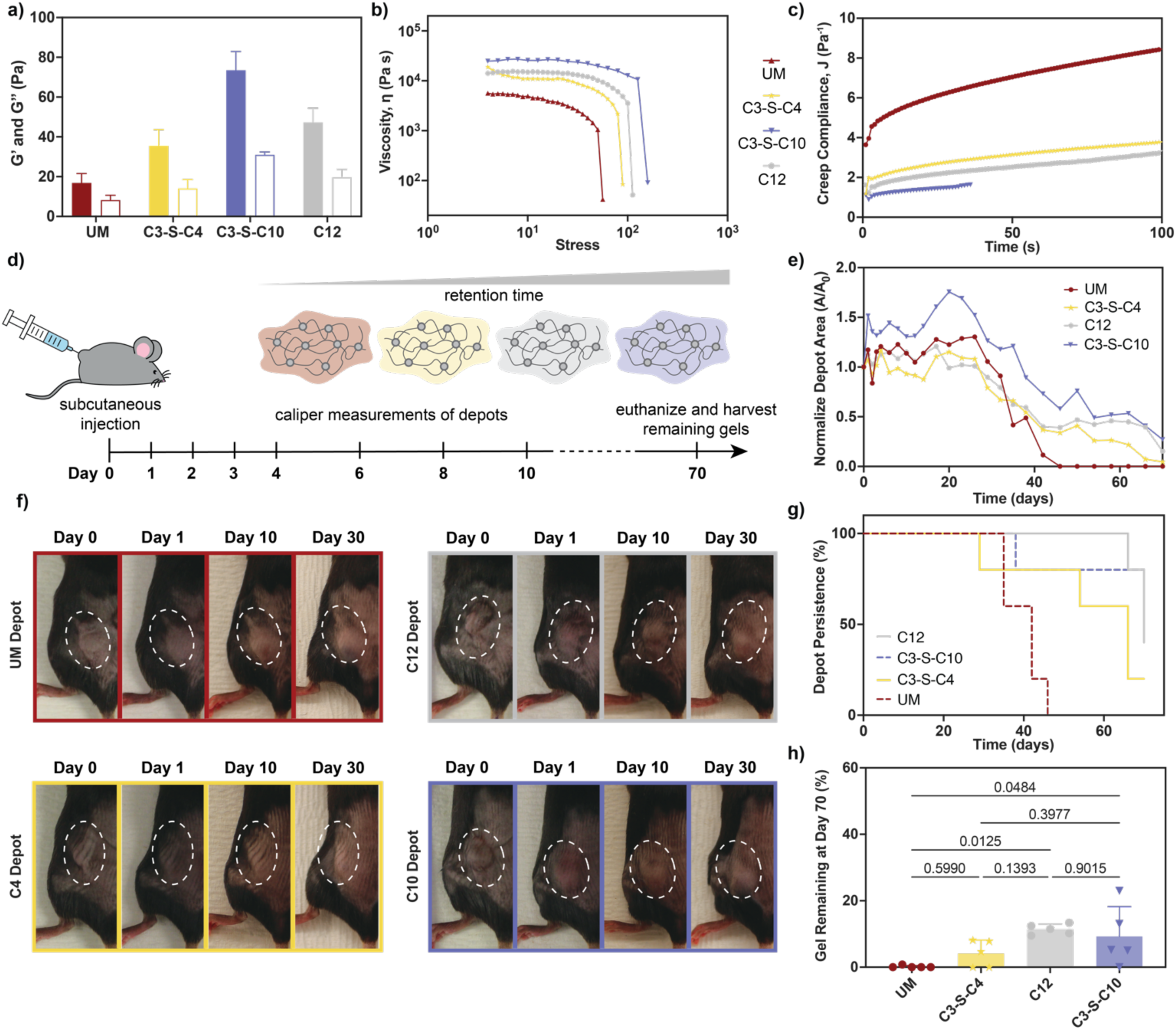
*In Vivo* Hydrogel Retention. a) Bar chart comparing average storage (G’, filled bars) and loss (G”, empty bars) moduli of PNP derivatives investigated *in vivo* (ω = 1 rad/s, 2% strain). Values reported are means ± SE of n = 3. b) Comparison of stress sweeps of PNP derivatives investigated *in vivo*. c) Comparison of creep compliance of PNP derivatives investigated *in vivo* when exposed to subcutaneously relevant 20 Pa stress. d) Timeline of *in vivo* experiment. e) Normalized observable depot areas over 70 days. f) Representative images of hydrogel depots of each PNP hydrogel derivative over 30 days. g) Fraction of mice with observable depots over 70 days. h) Gel mass remaining at day 70 measured from *in vivo* explants. Statistical analysis was run using GraphPad Prism using one-way ANOVA test with a Tukey’s multiple comparisons test across multiple groups. Values reported in plots d and h are means ± SE of n = 5.

## 3. Conclusions

Methods to gain control over mechanical properties and delivery timescales from PNP hydrogels can have critical impacts on therapeutic efficacy. For example, physical PNP hydrogel properties play a key role in the viability of adoptive cells^13^ and the magnitude of immune responses.^14,15,30^ We anticipate that this thiol–ene derivatization strategy will allow for greater control over mechanical properties and delivery profiles by specifically modifying the strength of polymer–nanoparticle interactions without altering hydrogel mesh size or component concentrations. Notably, by conducting a thorough investigation of rheological properties of our PNP hydrogel library, we have observed an interesting tradeoff between material retention time and recovery post-shear. Given that initial burst release from hydrogels after injection has remained an unmet problem,^28^ increasing recovery rates of materials post-injection might be a tunable handle to maintain drug exposure within the desired therapeutic window. Thus, future investigation into intermediate formulations that provide sustained release while offering more rapid recovery post-injection could reduce adverse effects associated with burst release.

Additionally, as previewed by our demonstration of RGD functionalization, this synthetic strategy is easily translatable to any cysteine-bearing peptide. Not only does this provide a route to easily functionalize with cell adhesive moieties that can improve cell biocompatibility, but it also opens exciting opportunities in the covalent attachment of peptides and small molecules with broad functions to the PNP scaffold. Small and hydrophilic cargo have traditionally been incompatible with delivery via PNP hydrogels as they readily diffuse out of the comparatively large mesh. Thus, strategies to incorporate small molecule agents have required burdensome conjugation to PEG-*b*-PLA nanoparticles.^22,31,32^ Thus, this HPMC modification strategy offers a new modular route to the delivery of thiol containing biologically active small molecules and peptides via PNP hydrogels.

In summary, we have presented a facile strategy for tuning PNP hydrogels based on facile thiol–ene click HPMC derivatization. This method takes advantage of the wealth of commercially available thiol containing compounds and cysteine-bearing peptides, greatly expanding the scope of previously accessible polymer–nanoparticle interactions that can be investigated. We used rheometry to probe the impact of HPMC pendant group identity on bulk material properties, and demonstrated that stiffness, flow and recovery behaviors could be readily tuned by changing HPMC pendant group identity. Finally, we showcase that these tunable mechanical properties could be leveraged to provide varied hydrogel retention times *in vivo*, highlighting the potential for this strategy for tailored therapeutic delivery.

## 4. Experimental

### 4.1 Materials

Hydroxypropylmethylcellulose 2910 (HPMC) was purchased from Sigma Aldrich (Product Number H3785). CGRGDSP peptide was custom ordered from Peptide 2.0. Unless otherwise noted, chemicals were purchased from Sigma Aldrich, TCI Chemicals or Fisher Scientific and used as received without further purification.

### 4.2 Polymer Characterization

All ^1^H NMR spectra were recorded on a Brucker Neo 500 MHz spectrometer and are reported relative to deuterated dimethyl sulfoxide (DMSO-D_6_, δ = 2.50 ppm) or chloroform (CDCl_3_, δ = 7.26 ppm). Average molecular weight and dispersity values were determined by size exclusion chromatography (SEC) eluting with *N,N*-dimethylformamide (0.01 M LiBr) and calibrated relative to poly(ethylene glycol) standards. Separation was done through two Jordi Labs Resolve Mixed Bed Low Divinylbenzene (DVB) columns in series and data was collected by a Dionex Ultimate 3000 Variable Wavelength detector and RefractoMax521 RI detector.

### 4.3 Synthesis of HPMC derivatives

#### HPMC-allyl Preparation

To an oven-dried round bottom flask containing stirred *N*-Methyl-2-pyrrolidone (NMP) (280 mL, anhydrous) was slowly added HPMC (7 g, ~30 mmol cellulose repeat units). The solution was allowed to stir at room temperature under inert atmosphere until HPMC had fully dissolved. The solution was then heated to 50 °C before adding *N,N*-Diisopropylamine (0.89 mL, 5.09 mmol, 0.17 eq/repeat) dropwise. In a separate vial, allyl isocyanate (0.82 mL, 3.6 mmol, 0.12 eq/repeat) was pre-mixed with NMP (10 mL NMP) before it was added dropwise to the stirred HPMC solution. Once added, the reaction was allowed to cool to room temperature and stir under nitrogen atmosphere for and additional 24 hours. The material was isolated by precipitation into a stirred mixture of acetone/methanol (70:30, 2 L). The isolated solid was then washed thoroughly with acetone before it was dried under reduced pressure. For thiol–ene click reactions, this HPMC-allyl was taken forward without additional purification. HPMC-allyl used for subsequent rheological, *in vitro* and *in vivo* experiments was further purified by dialysis against milli-Q water (MWCO = 3.5 kDa) for 2 days followed by lyophilization. Allyl incorporation was determined to be 4% by ^1^H NMR.

#### General Procedure for Thiol-ene click

To a 20-mL scintillation vial equipped with stir bar, HPMC-allyl (300 mg, ~1.2 mmol cellulose repeat units, ~0.14 mmol ene) and the desired thiol (7.5 eq/ene) were dissolved in NMP (12 mL). A stock solution of 2,2-Dimethoxy-2-phenylacetophenone in NMP (185 mg/mL) was then added to the vial (0.1 mL, 0.07 mmol, 0.5 eq/ene) before irradiating in an LZC-4V photoreactor at 355 nm (4 mW/cm^2^) for 90 minutes. After irradiation the material was isolated by precipitation into acetone (200 mL). The isolated solid was then washed thoroughly with acetone before it was dissolved in milli-Q water (30 mL) and transfered into hydrated 3.5 kDa MWCO dialysis tubing. The polymer was purified by dialysis against milli-Q water (1 L) for 2 days followed by lyophilization. Quantitative conversion of allyl groups was confirmed by ^1^H NMR.

### 4.4 Synthesis of poly(ethylene glycol)-*b*-poly(lactic acid) (PEG-*b*-PLA)

PEG-*b*-PLA was prepared by ring opening polymerization of D,L-lactide from poly(ethylene glycol) monomethyl ether (PEG-OMe) (5 kDa, Sigma Aldrich) similar to previous reports.^12,16,25^ To ensure exclusion of water, the following purifications were conducted prior to polymerization: 1,8- diazabicyclo[5.4.0]undec-7-ene (DBU) was purified by distillation under reduced pressure and stored over molecular sieves until use; D,L-lactide was recrystallized 3 times from anhydrous ethyl acetate; PEG-OMe was dried at 100 °C under vacuum for 3 hours directly before use; and dichloromethane (DCM) was dispensed from a solvent purification system immediately before use. In an oven-dried round bottom flask under nitrogen atmosphere, D,L-lactide (8 g, 55.5 mmmol, 139 eq) was dissolved in DCM (36 mL). Once dissolved, PEG-OMe (2 g, 0.4 mmol, 1 eq) dissolved in DCM (4 mL) was added. To initiate the polymerization a stock solution of DBU (150 uL/mL DCM) was quickly added (400 uL stock, 0.4 mmol, 1 eq) and the polymerization was conducted for 8 minutes. The polymerization was then terminated by addition of 0.5 mL stock solution of acetic acid in acetone (3-4 drops/mL). Conversion was determined to be 90 % (*M_n,theor_* = 23,000 Da) by ^1^H NMR. The polymer was purified by precipitation into a 1:1 mixture of hexanes/diethyl ether (400 mL). Molecular weight was determined by ^1^H NMR: *M_n,NMR_* = 24,600 Da and GPC (DMF) *M_n,GPC_* = 21,000 Da, Đ = 1.09.

### 4.5 Nanoparticle Preparation and Characterization

PEG-*b*-PLA NPs were prepared within 24 hours of use by nanoprecipitation as previously reported.^12,16,25^ A solution of PEG-*b*-PLA (50 mg/mL) was prepared in a mixture of acetonitrile and dimethyl sulfoxide (75:25). For each batch of nanoparticles, 1 mL of this PEG-b-PLA solution was added dropwise into 10 mL milli-Q water stirred at 600 rpm. Nanoparticles were then concentrated by centrifugal filtration (MWCO 10 kDa) and resuspended in PBS to afford a 20 wt% stock solution. Hydrodynamic diameter and dispersity of nanoparticle batches were assessed by dynamic light scattering (DynaPro II plate reader, Wyatt Technology): *D_H_* = 28 – 32 nm, *PDI* < 15% for all batches.

### 4.6 General PNP Hydrogel Preparation

PNP hydrogels were prepared similar to previous reports.^12,16,25^ Stock solutions of each HPMC-X were prepared by dissolving at 6 wt% in PBS over several days on a nutator with additional agitation introduced as needed to ensure sufficient dissolution. Once dissolved, stock solutions were stored at 4 °C until use. For rheological experiments, hydrogels (600 uL) were prepared by mixing 6 wt% HPMC-X stock solutions (100 mg), 20 wt% PEG-*b*-PLA NPs stock solution (150 uL) and PBS (350 uL) in Eppendorf tubes to afford final concentrations of 1 wt% HPMC-X and 5 wt% PEG-*b*-PLA. Gels were periodically placed on a benchtop centrifuge to remove bubbles during mixing. For *in vitro* and *in vivo* experiments, PEG-*b*-PLA NPs stock solution, PBS, and dyes were loaded into one syringe, HPMC-X was loaded into a second syringe, and the components were mixed using an elbow connector. After mixing, the elbow mixer was replaced with a needle for injection into capillary tubes or mice.

### 4.7 Rheology

Rheological experiments on PNP hydrogels were conducted at 25 °C using a 20 mm diameter serrated parallel plate at a 500 µm gap on a stress-controlled TA Instruments DHR-2 rheometer equipped with solvent trap. For 1 wt% HPMC-X solutions, testing was conducted using a 40 mm, 2.018 ° cone and plate geometry. Frequency sweeps for all PNP hydrogels were performed at a strain of 1% within the linear viscoelastic regime. Flow sweeps were performed from high to low shear rates with steady state sensing. Stress controlled flow measurements were performed from low to high stress with steady state sensing and 10 or 20 points per decade. For stress relaxation experiments, a 5% step strain was applied, and the corresponding stress and modulus was measured for 1000 s. Relaxation times were determined by fitting to Kohlrausch’s stretched-exponential relaxation model^33^ in GraphPad Prism (**Supplemental Figure 20**). Creep experiments measured strain rate at fixed stress. An initial stress of 0.5 Pa was applied to the material for 20 s. The desired stress (5–200 Pa) was then applied, and corresponding strain percent was measured for 300 s. Strain rate at a given stress was obtained by fitting a slope for the linear region of the strain vs time curve in GraphPad Prism. Stress overshoot experiments were conducted similar to previously described.^16^ For each step, a flow step was applied (10 s^−1^) followed by a specified wait time (2, 4, 6, 10, 12, 20, 40, 60 or 120 s) before moving onto the next step (**Figure 4b**). Characteristic relaxation times were determined by fitting the stress overshoot as a function of wait time to an exponential plateau in GraphPad Prism (**Supplemental Figures S25-26**).

### 4.8 Fluorescence Recovery after Photobleaching (FRAP) Analysis

FRAP analysis was conducted as previously described.^15^ PNP hydrogels containing 0.5 mg/mL BSA-FITC (Sigma) were prepared, placed on glass slides and imaged using a confocal ZEISS LSM780 microscope (n=3 per hydrogel type). Samples were imaged to obtain an initial level of fluorescence (excitation = 488 nm) before a high intensity laser (488 nm) was used to bleach a region of interest (ROI) with 12.5 μm diameter for 10 s. Fluorescence data was then recorded for 200 seconds to obtain an exponential fluorescence recovery curve. Diffusivity values were calculated according to,

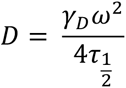

Where ω is the radius of the bleached ROI and the constant γ_D_ is calculated as τ_1/2_/τ_D_. Both τ_1/2_, the half-time of recovery, and τ_D,_ the characteristic recovery time, are given by the ZEN software.

The diffusivity of BSA in PBS was calculated according to the Stokes-Einstein Equation,

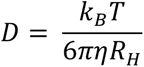

Where *k*_B_ is Boltzmann’s constant, *T* is temperature in Kelvin, *η* is solvent viscosity (0.8872 mPa.s for PBS at 25 °C),^34^ and *R*_H_ is the hydrodynamic radius (3.4 nm for BSA).^35^

### 4.9 *In Vitro* Capillary Release

Glass capillary tubes (inner diameter = 3 mm) were plugged at one end with epoxy before 100 uL of each PNP hydrogel containing 25 ug BSA-Alexa 647 (Invitrogen) was injected (n=3). Hydrogels were carefully topped with 200 uL of PBS and capillaries were sealed with parafilm and incubated at 37 °C. At each time point, the PBS was removed and replaced with fresh PBS. Release into PBS was determined by fluorescent measurement (excitation = 650 nm, emission = 668 nm) relative to calibration curve of known BSA-Alexa 647 concentrations.

### 4.10 *In Vivo* Methodology

Seven-week-old female C57BL/6 mice were purchased from Charles River and housed in the animal facility at Stanford University. Mice were cared for following the Institutional Animal Care and Use guidelines. All animal studies were performed in accordance with the National Institutes of Health guidelines and the approval of Stanford Administrative Panel on Laboratory Animal Care (protocol APLAC-32109). Mice were shaved on both flanks prior to injections. Each mouse received two subcutaneous injections (100 μL) of PNP hydrogel on either flank while under brief isoflurane anesthesia. Depots were measured with calipers in the x and y plane at each timepoint and depot areas were estimated as an ellipse. After 70 days, mice were euthanized and remaining PNP gels were excised and weighed to determined remaining mass percentage relative to initial injection.

## 5. Author Contributions

S.J.B., N.E., and E.A.A conceived the ideas. S.J.B., N.E., and E.A.A. designed the experiments. S.J.B., N.E., E.B., and C.K.J. conducted the experiments. S.J.B., N.E., and S.S. analyzed the data. S.J.B. wrote the paper. S.J.B., N.E., and E.A.A. revised the paper. All authors have given approval to the final version of the manuscript.

## Acknowledgements

The research reported here was financially supported by the Bill & Melinda Gates Foundation (INV-027411). The instrument used for ^1^H NMR data acquisition was funded significantly by an NIH High End Instrumentation grant (1 S10 OD028697-01). S.J.B. would like to thank Stanford’s Office for Postdoctoral Affairs for support through a PRISM Baker Fellowship. N.E. was supported by an NSF Graduate Research Fellowship (DGE-2146755). C.J. was supported by an NSF Graduate Research Fellowship. The authors would also like to thank Prof. Sarah Heilshorn for use of a Leica DMi8 microscope for cellular imaging.

## 7. Supporting Information

HPMC ^1^H NMRs, cell encapsulation studies, supplemental rheological data and photographs of *in vivo* excisions can be found in the Supporting Information.

