## Supporting Information for "A thiol–ene click-based strategy to customize injectable polymer–nanoparticle hydrogel properties for therapeutic delivery"

### 1. Characterization of HPMC derivatives

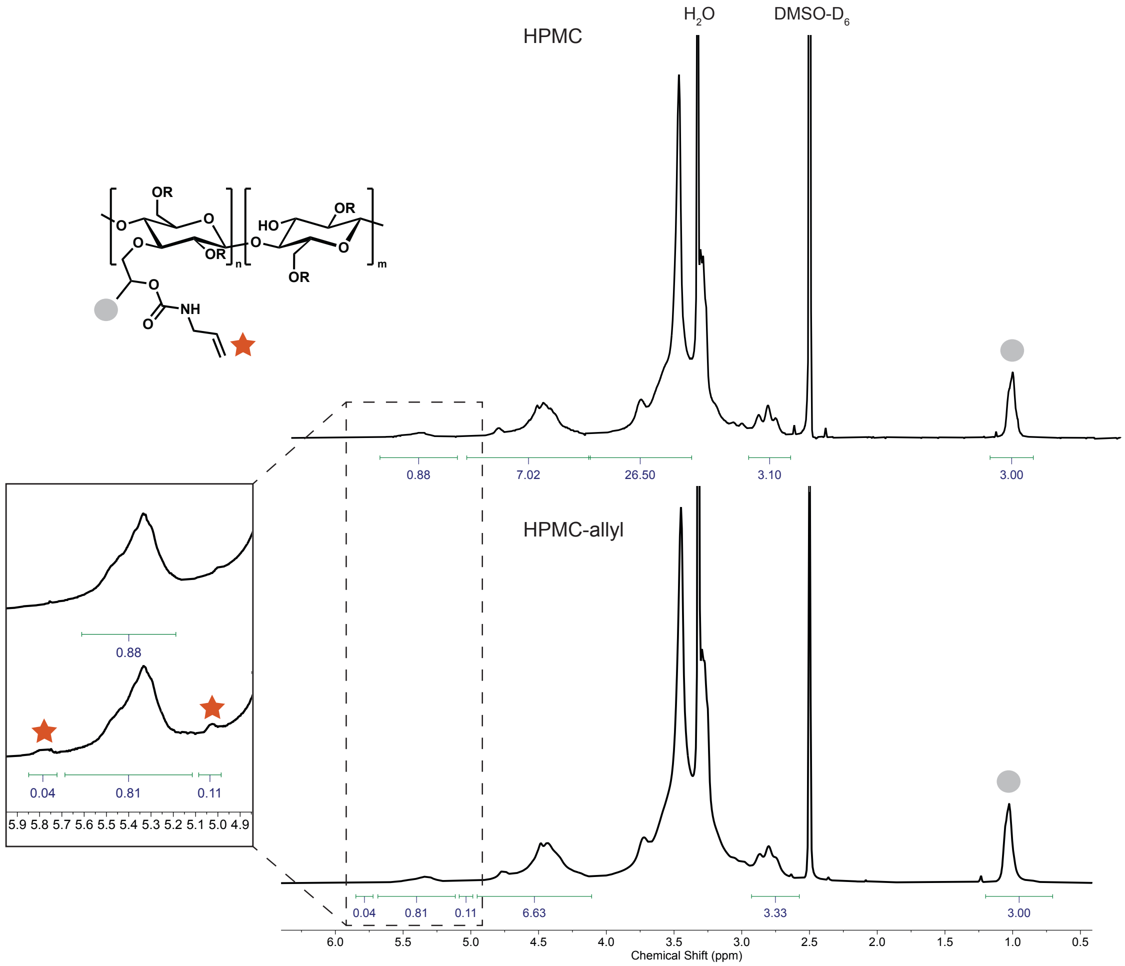

**Figure S 1.** Overlay of $^{1}H$ NMRs of commercial HPMC (top) and isolated product (bottom) after functionalization with allyl isocyanate. Integrations indicate 5% functionalization relative to hydroxypropyl protons at 1.0 ppm.

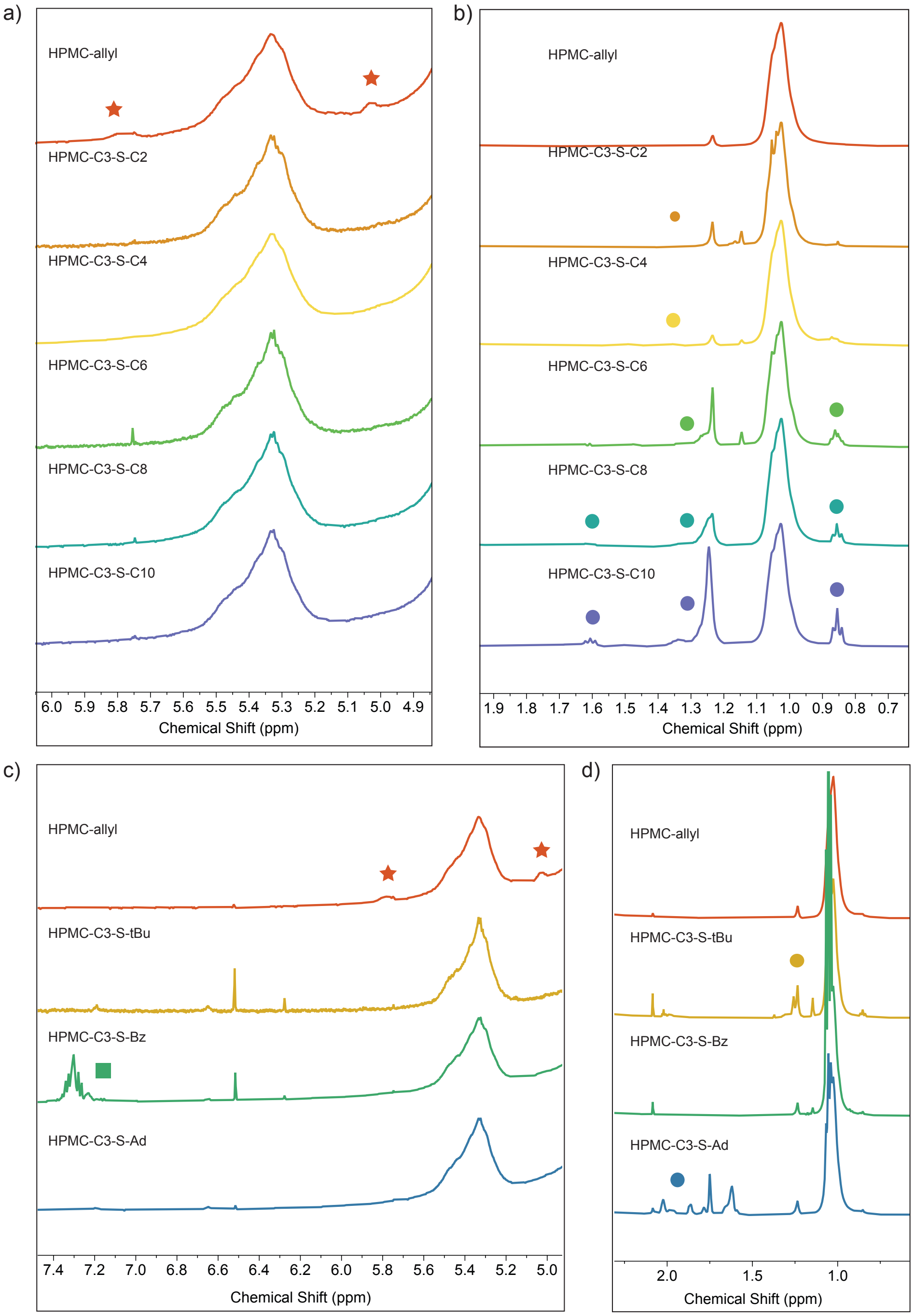

**Figure S 2.** Overlay $^1$H NMR spectra of HPMC-allyl with products obtained from thiol–ene clicks. Consumption of allylic protons at 5.0 and 5.8 ppm (orange stars) observed after clicking to a) linear and c) steric/pi stacking thiols. Appearance of diagnostic alkyl protons observed at 0.8 – 2.0 ppm (circles) observed after clicking to b) linear and d) sterically hindered thiols; and c) aromatic protons at 7.3 ppm (green square) after clicking to Bz thiol.

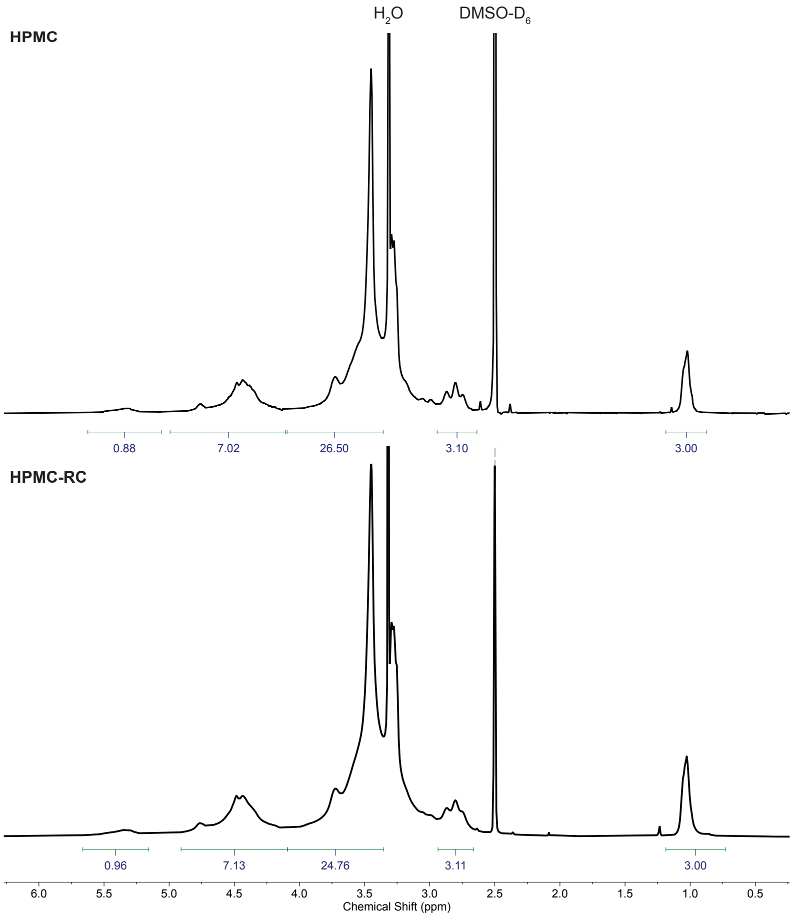

**Figure S 3.** Overlay of $^1$H NMRs of commercial HPMC (top) and control material exposed to radical control conditions (HPMC-RC) indicates presence of free radicals does not lead to side reactions with the HPMC.

### 2. Cell Encapsulated RGD PNP Hydrogels

10x incorporation HPMC-allyl parent material was first prepared following the protocol provided in the main text **Section 4.3** using 1.2 equivalents allyl isocyanate/cellulose repeat. The general procedure for thiol–ene click was then followed using 2.5 equivalents of CGRGDSP peptide/ene. $^1$H NMR indicated 15% functionalization relative to hydroxypropyl group, which was used to determine RGD concentration in final hydrogels.

NIH/3T3 fibroblasts were purchased from ATCC and cultured at 37 °C with 5% $CO_2$ in Dulbecco's Modified Eagle Medium (DMEM) supplemented with 10% fetal bovine serum and 1% antibiotic (100 units/mL penicillin, 100 ug/mL streptomycin). Cells were stained by incubating in 10 µM CellTracker™ Green BODIPY™ Dye (Invitrogen) in serum-free media for 45 minutes before trypsinization. PNP hydrogels were prepared following the general procedure by mixing in syringes with an elbow mixer. Cell stock and additional culturing media was added to the nanoparticle stock solution before mixing with HPMC stock solutions to provide a final gel concentrations of 5e6 cells/mL, 5 wt% PEG-*b*-PLA NPs, 1 wt% HPMC-C12 and either 0 or 0.2 wt% HPMC-RGD (corresponding to 0 or 1.2 mM RGD, respectively). After mixing, 20 µL gels were dispensed into a 12-well plate and 300 uL media was added to each gel. Before imaging, excess DMEM was removed, and coverslips were placed atop each gel.

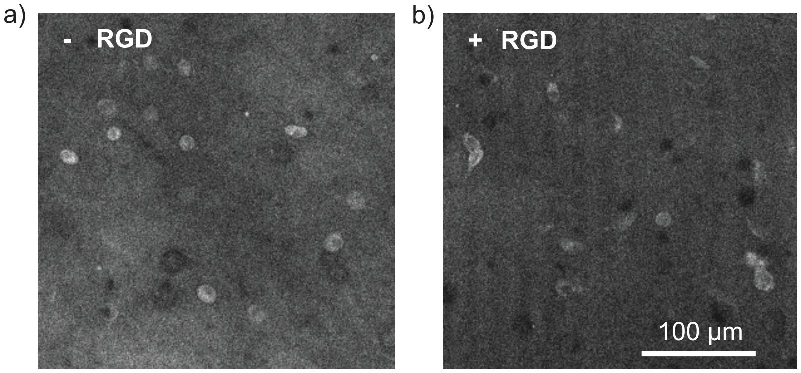

**Figure S 4.** Fluorescence imaging using Leica DMi8 microscope (excitation = 522 nm, emission = 529 nm) of cells 7 hours after being encapsulated in a) non-RGD containing PNP hydrogels and b) PNP hydrogels containing HPMC-RGD.

### 3. Supplemental Rheological Data

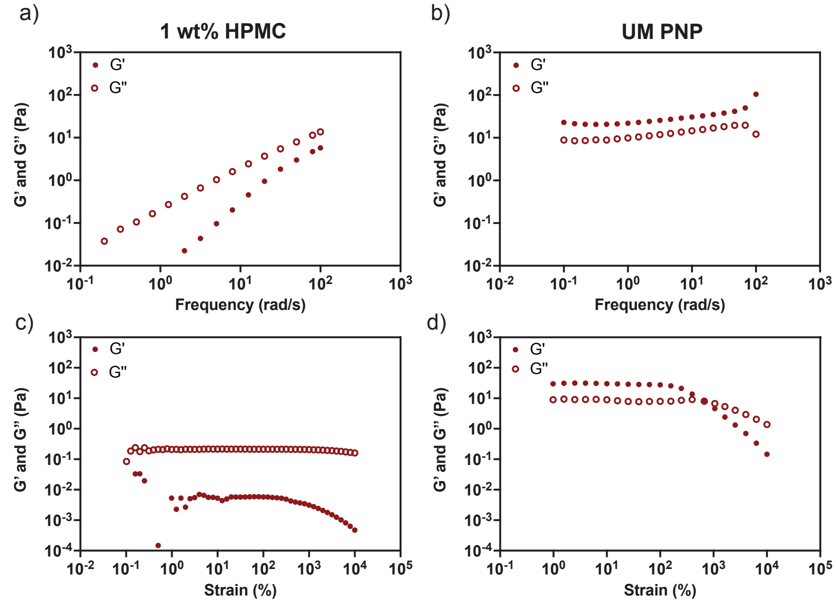

**Figure S 5.** Frequency sweeps of a) 1 wt% solution of HPMC compared to b) mixture of 1 wt% solution of HPMC with 5 wt% PEG-*b*-PLA NPs at 1% strain. c) Amplitude sweep of 1 wt% solution of HPMC at ω = 0.2 rad/s. d) Amplitude sweep of the mixture of 1 wt% solution of HPMC with 5 wt% PEG-*b*-PLA NPs at ω = 1 rad/s.

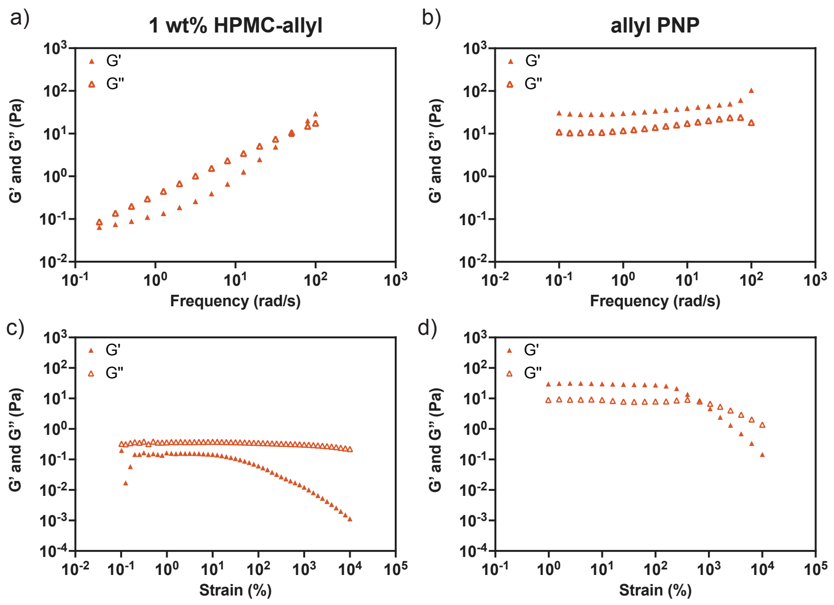

**Figure S 6.** Frequency sweeps of a) 1 wt% solution of HPMC-allyl compared to b) mixture of 1 wt% solution of HPMC-allyl with 5 wt% PEG-*b*-PLA NPs at 1% strain. c) Amplitude sweep of 1 wt% solution of HPMC-allyl at ω = 0.2 rad/s. d) Amplitude sweep of the mixture of 1 wt% solution of HPMC-allyl with 5 wt% PEG-*b*-PLA NPs at ω = 1 rad/s.

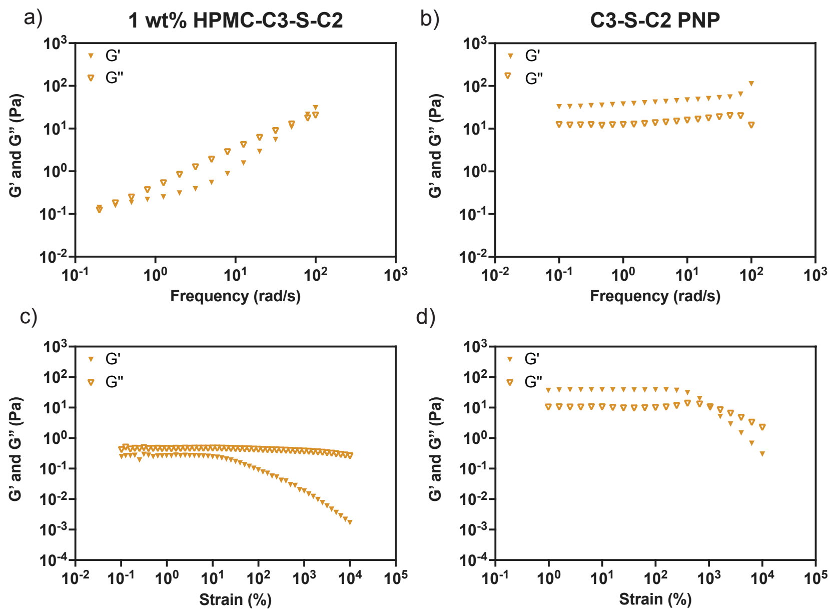

**Figure S 7.** Frequency sweeps of a) 1 wt% solution of HPMC-C3-S-C2 compared to b) mixture of 1 wt% solution of HPMC-C3-S-C2 with 5 wt% PEG-*b*-PLA NPs at 1% strain. c) Amplitude sweep of 1 wt% solution of HPMC-C3-S-C2 at ω = 0.2 rad/s. d) Amplitude sweep of the mixture of 1 wt% solution of HPMC-C3-S-C2 with 5 wt% PEG-*b*-PLA NPs at ω = 1 rad/s.

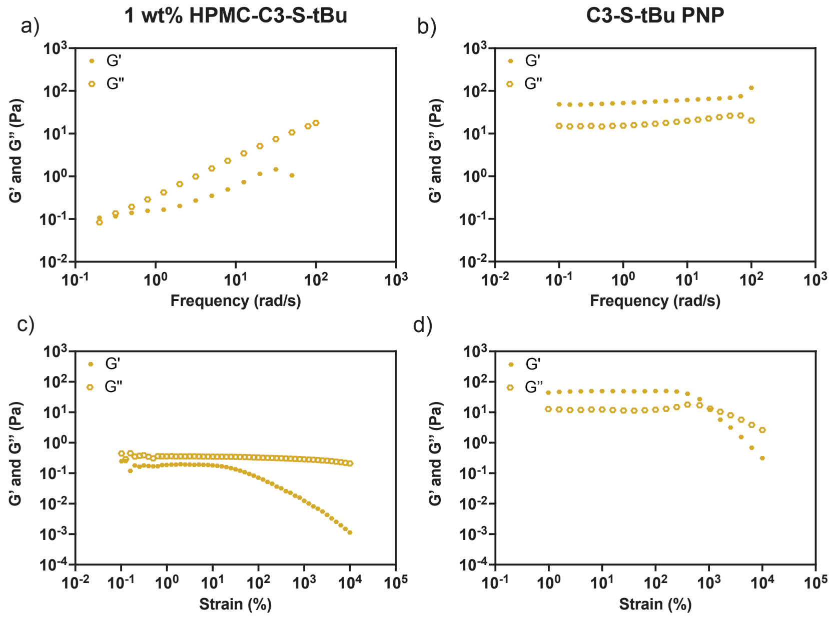

**Figure S 8.** Frequency sweeps of a) 1 wt% solution of HPMC-C3-S-tBu compared to b) mixture of 1 wt% solution of HPMC-C3-S-tBu with 5 wt% PEG-*b*-PLA NPs at 1% strain. c) Amplitude sweep of 1 wt% solution of HPMC-C3-S-tBu at ω = 0.2 rad/s. d) Amplitude sweep of the mixture of 1 wt% solution of HPMC-C3-S-tBu with 5 wt% PEG-*b*-PLA NPs at ω = 1 rad/s.

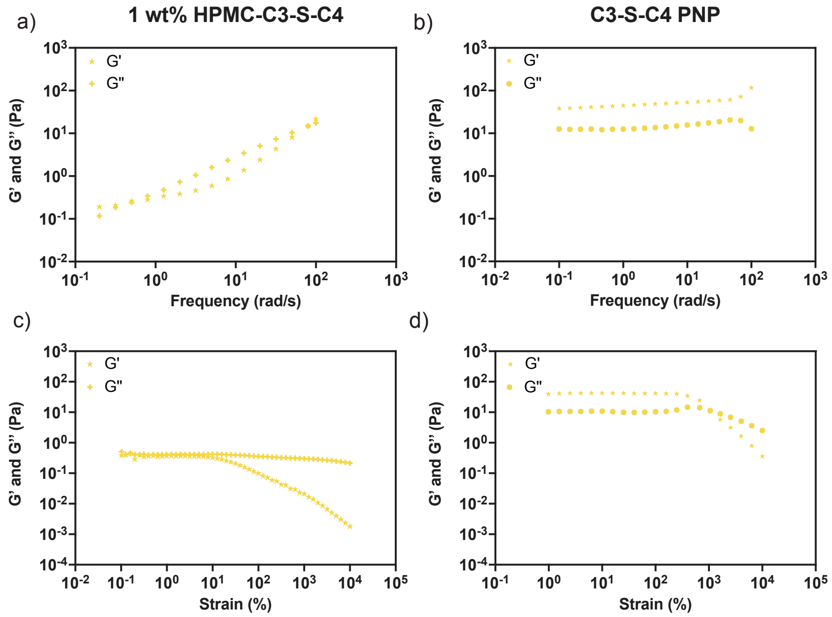

**Figure S 9.** Frequency sweeps of a) 1 wt% solution of HPMC-C3-S-C4 compared to b) mixture of 1 wt% solution of HPMC-C3-S-C4 with 5 wt% PEG-*b*-PLA NPs at 1% strain. c) Amplitude sweep of 1 wt% solution of HPMC-C3-S-C4 at ω = 0.2 rad/s. d) Amplitude sweep of the mixture of 1 wt% solution of HPMC-C3-S-C4 with 5 wt% PEG-*b*-PLA NPs at ω = 1 rad/s.

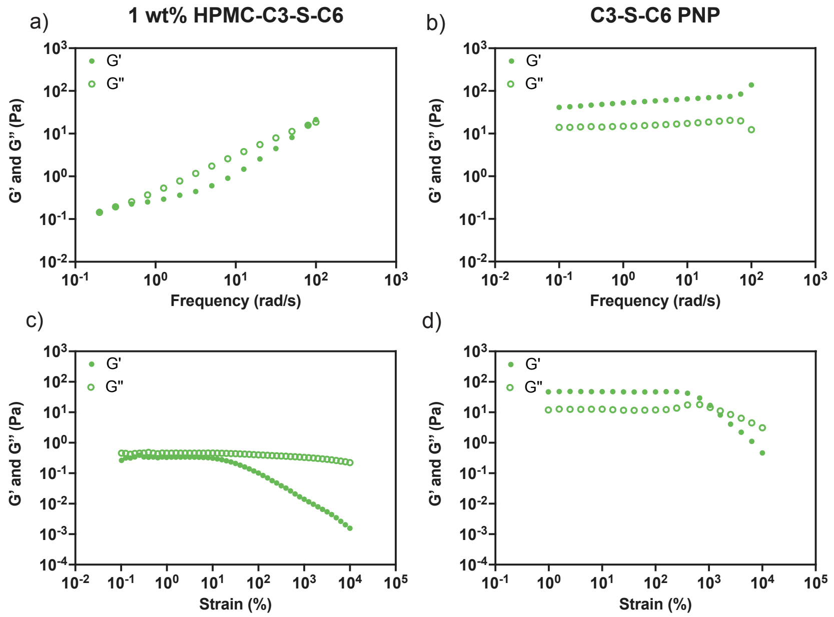

**Figure S 10.** Frequency sweeps of a) 1 wt% solution of HPMC-C3-S-C6 compared to b) mixture of 1 wt% solution of HPMC-C3-S-C6 with 5 wt% PEG-*b*-PLA NPs at 1% strain. c) Amplitude sweep of 1 wt% solution of HPMC-C3-S-C6 at ω = 0.2 rad/s. d) Amplitude sweep of the mixture of 1 wt% solution of HPMC-C3-S-C6 with 5 wt% PEG-*b*-PLA NPs at ω = 1 rad/s.

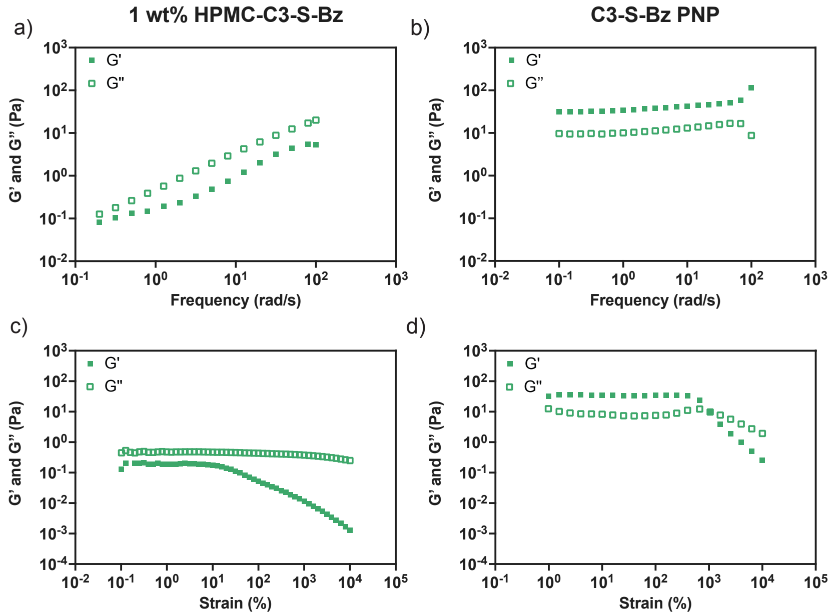

**Figure S 11.** Frequency sweeps of a) 1 wt% solution of HPMC-C3-S-Bz compared to b) mixture of 1 wt% solution of HPMC-C3-S-Bz with 5 wt% PEG-*b*-PLA NPs at 1% strain. c) Amplitude sweep of 1 wt% solution of HPMC-C3-S-Bz at ω = 0.2 rad/s. d) Amplitude sweep of the mixture of 1 wt% solution of HPMC-C3-S-Bz with 5 wt% PEG-*b*-PLA NPs at ω = 1 rad/s.

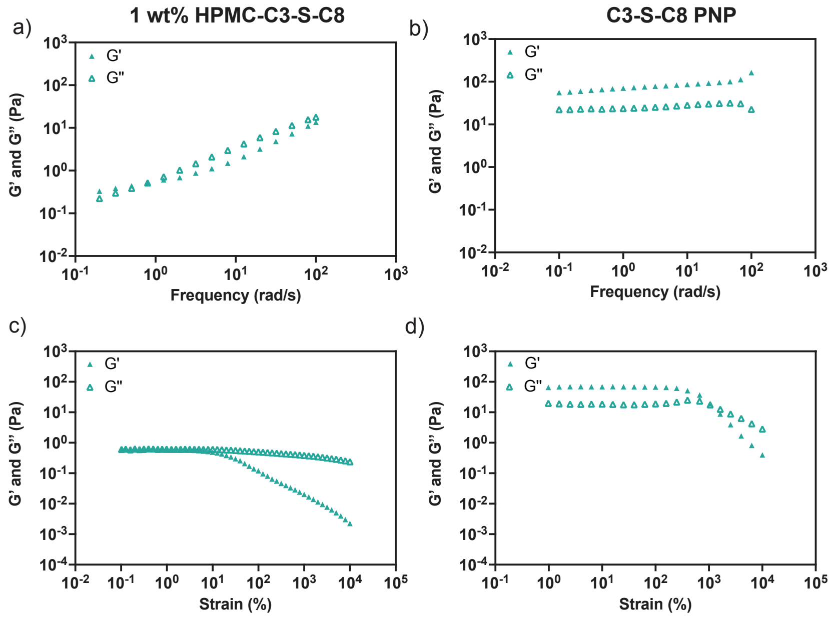

**Figure S 12.** Frequency sweeps of a) 1 wt% solution of HPMC-C3-S-C8 compared to b) mixture of 1 wt% solution of HPMC-C3-S-C8 with 5 wt% PEG-*b*-PLA NPs at 1% strain. c) Amplitude sweep of 1 wt% solution of HPMC-C3-S-C8 at ω = 0.2 rad/s. d) Amplitude sweep of the mixture of 1 wt% solution of HPMC-C3-S-C8 with 5 wt% PEG-*b*-PLA NPs at ω = 1 rad/s.

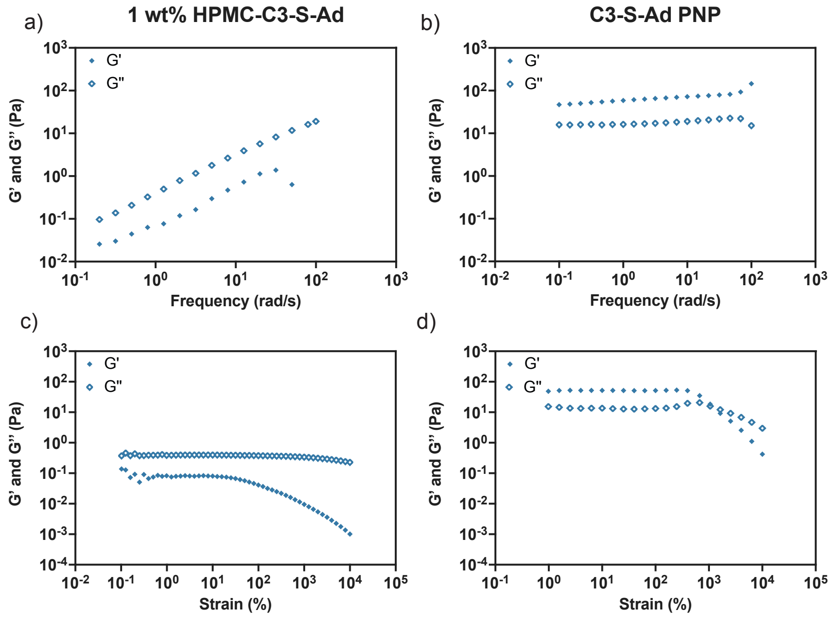

**Figure S 13.** Frequency sweeps of a) 1 wt% solution of HPMC-C3-S-Ad compared to b) mixture of 1 wt% solution of HPMC-C3-S-Ad with 5 wt% PEG-*b*-PLA NPs at 1% strain. c) Amplitude sweep of 1 wt% solution of HPMC-C3-S-Ad at ω = 0.2 rad/s. d) Amplitude sweep of the mixture of 1 wt% solution of HPMC-C3-S-Ad with 5 wt% PEG-*b*-PLA NPs at ω = 1 rad/s.

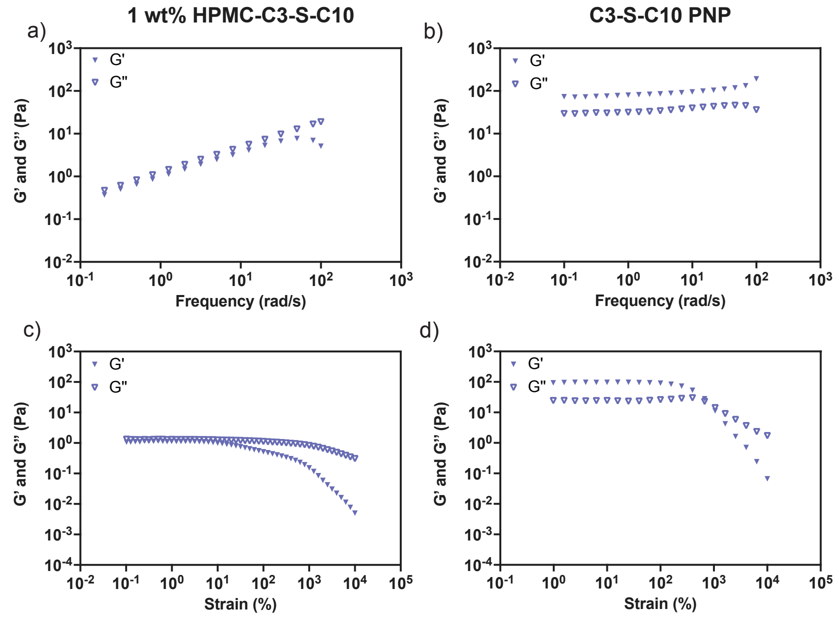

**Figure S 14.** Frequency sweeps of a) 1 wt% solution of HPMC-C3-S-C10 compared to b) mixture of 1 wt% solution of HPMC-C3-S-C10 with 5 wt% PEG-*b*-PLA NPs at 1% strain. c) Amplitude sweep of 1 wt% solution of HPMC-C3-S-C10 at ω = 0.2 rad/s. d) Amplitude sweep of the mixture of 1 wt% solution of HPMC-C3-S-C10 with 5 wt% PEG-*b*-PLA NPs at ω = 1 rad/s.

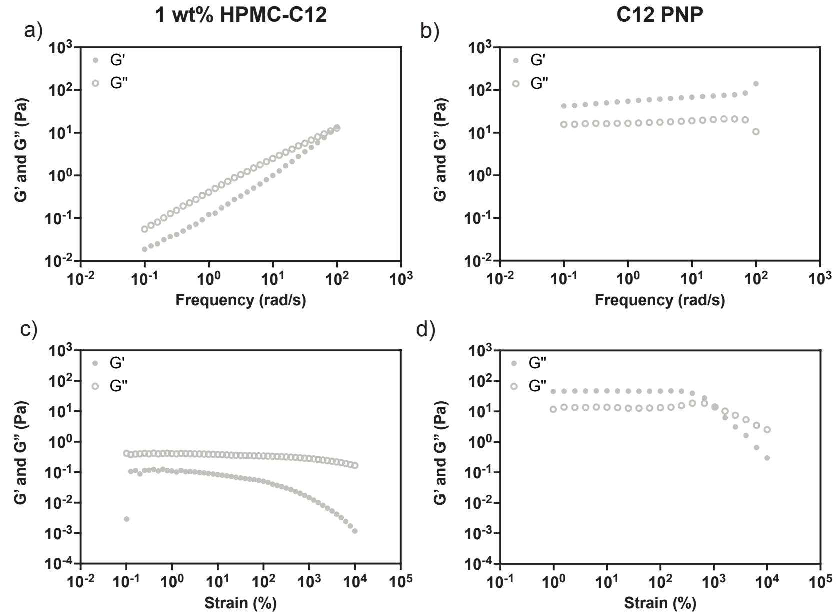

**Figure S 15.** Frequency sweeps of a) 1 wt% solution of HPMC-C12 compared to b) mixture of 1 wt% solution of HPMC-C12 with 5 wt% PEG-*b*-PLA NPs at 1% strain. c) Amplitude sweep of 1 wt% solution of HPMC-C12 at ω = 0.2 rad/s. d) Amplitude sweep of the mixture of 1 wt% solution of HPMC-C12 with 5 wt% PEG-*b*-PLA NPs at ω = 1 rad/s.

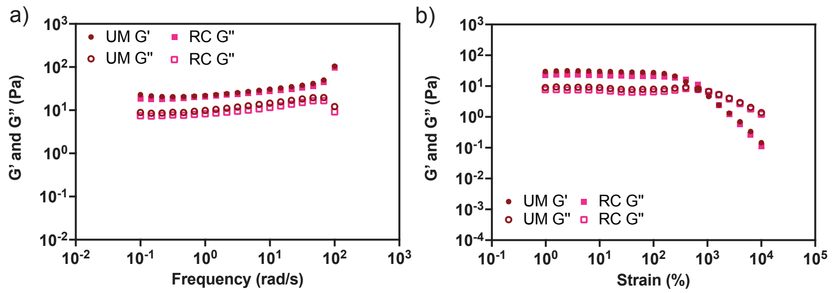

**Figure S 16.** a) Frequency sweeps (1% strain) and b) amplitude sweeps (ω = 1 rad/s) comparing properties of PNPs prepared with commercial HPMC (UM) and with HPMC exposed to radical control conditions (RC) indicate minimal differences occur in HPMC properties following exposure to free radical conditions during thiol–ene click reaction.

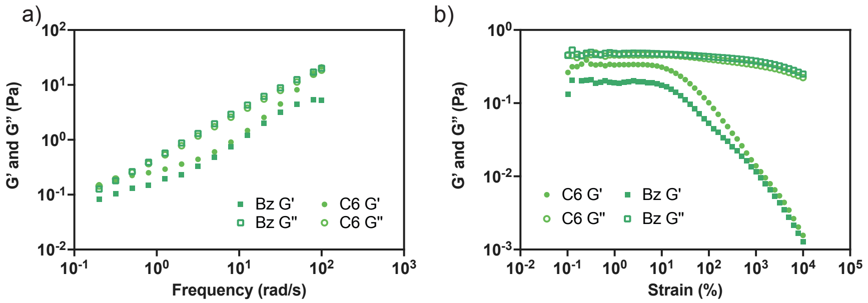

**Figure S 17.** a) Frequency sweeps (1% strain) and b) amplitude sweeps (ω = 0.2 rad/s) comparing properties of 1 wt% solution of HPMC-C6 and HPMC-Bz.

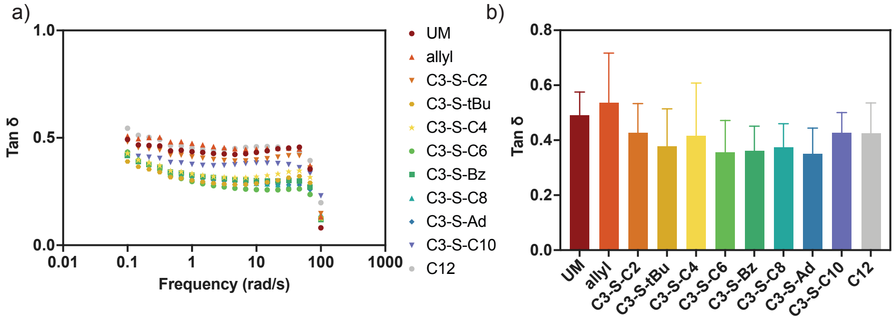

**Figure S 18.** a) Representative frequency sweep plotting tan δ for each PNP derivative (2% strain). b) Bar chart representing average tan δ of three independent runs at ω = 1 rad/s.

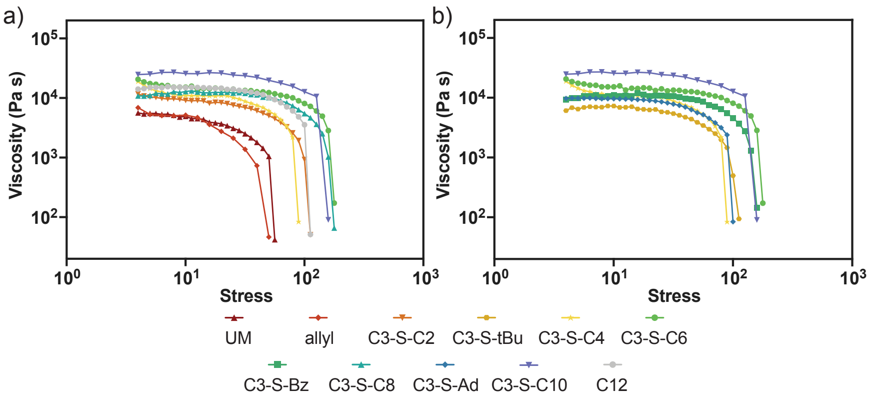

**Figure S 19.** Stress controlled flow measurements for PNP hydrogel derivatives.

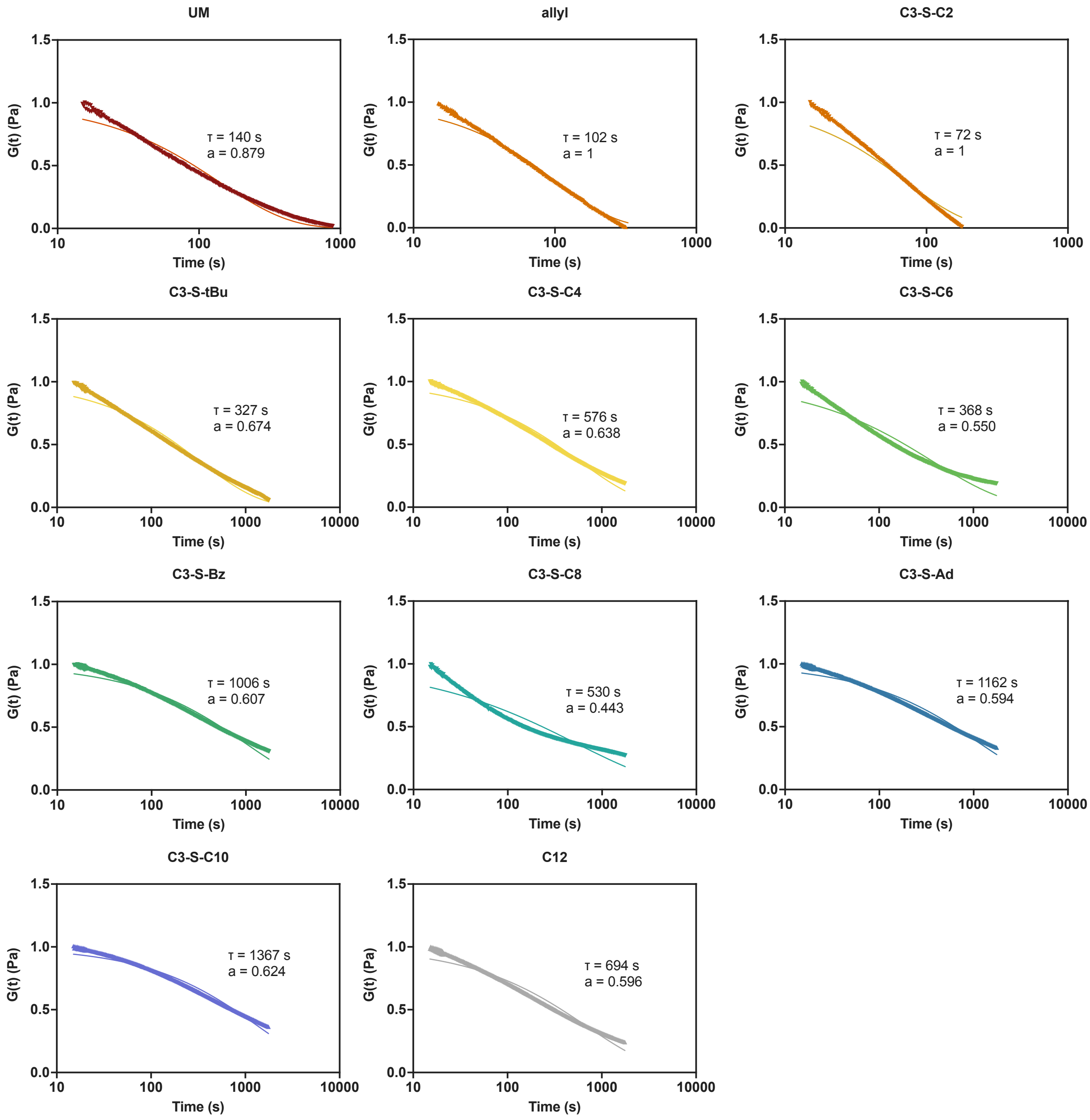

**Figure S 20.** Fitting of stress relaxation data to Kohlrausch's stretched-exponential model, $\frac{G}{G_0} = \exp(-\left(\frac{t}{\tau_R}\right)^a)$, to obtain characteristic relaxation times $\tau_R$.

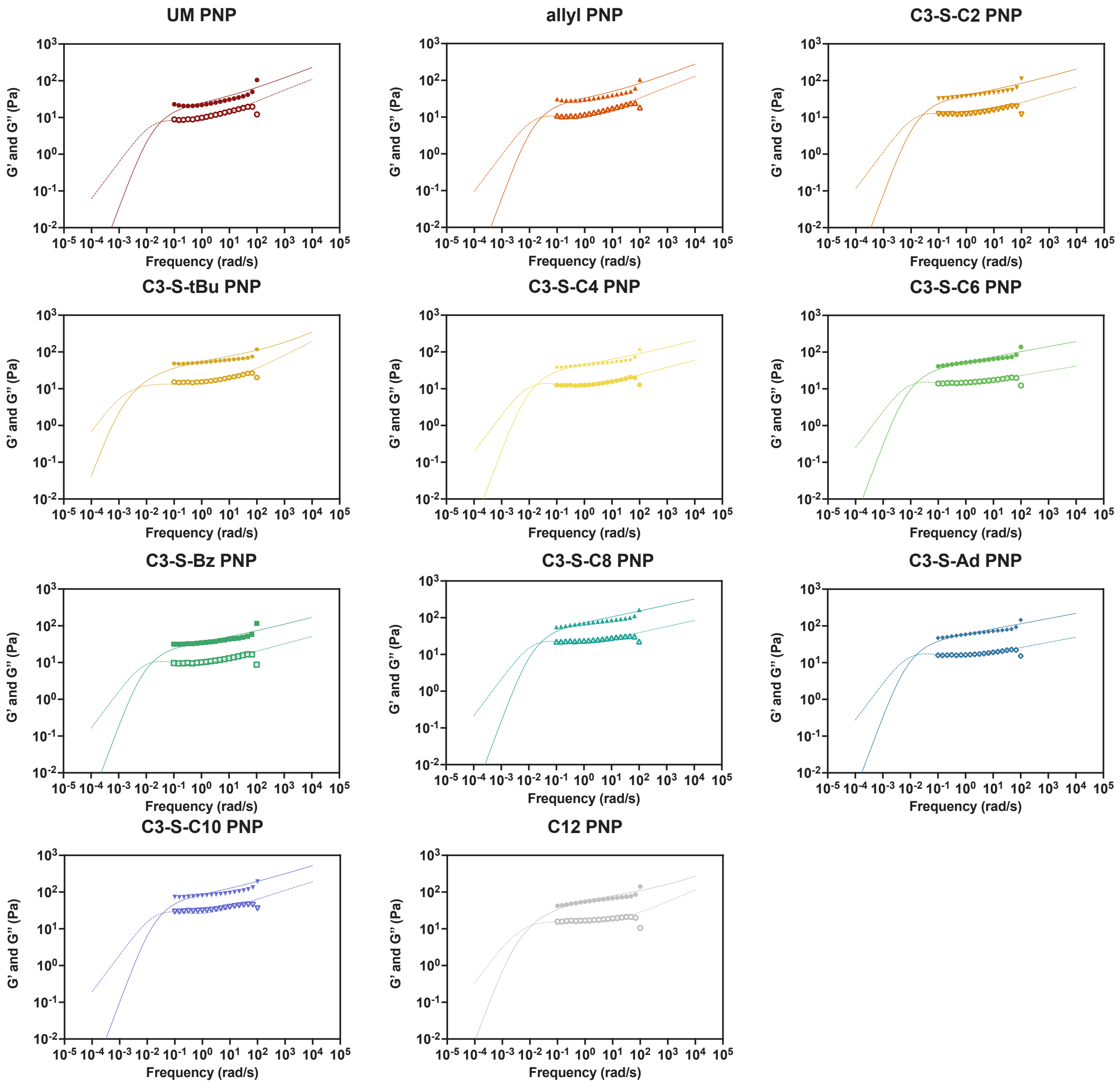

**Figure S 21.** Fitting of frequency sweeps to continuous viscoelastic relaxation spectra. Frequency sweep data were fit to the 7-parameter modified BSW distribution within the continuous spectrum framework, containing a viscoelastic fluid and a glassy contribution. The relaxation times were computed as the first mean timescales of the viscoelastic fluid component of the distribution obtained for each hydrogel system after the fit.

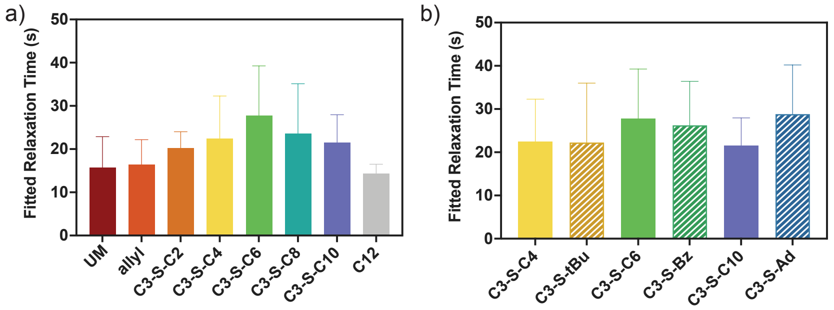

**Figure S 22.** Bar charts comparing relaxation times a) increasing pendant chain length and b) steric effects determined by fits from Figure SX with n=3 independent runs per hydrogel type.

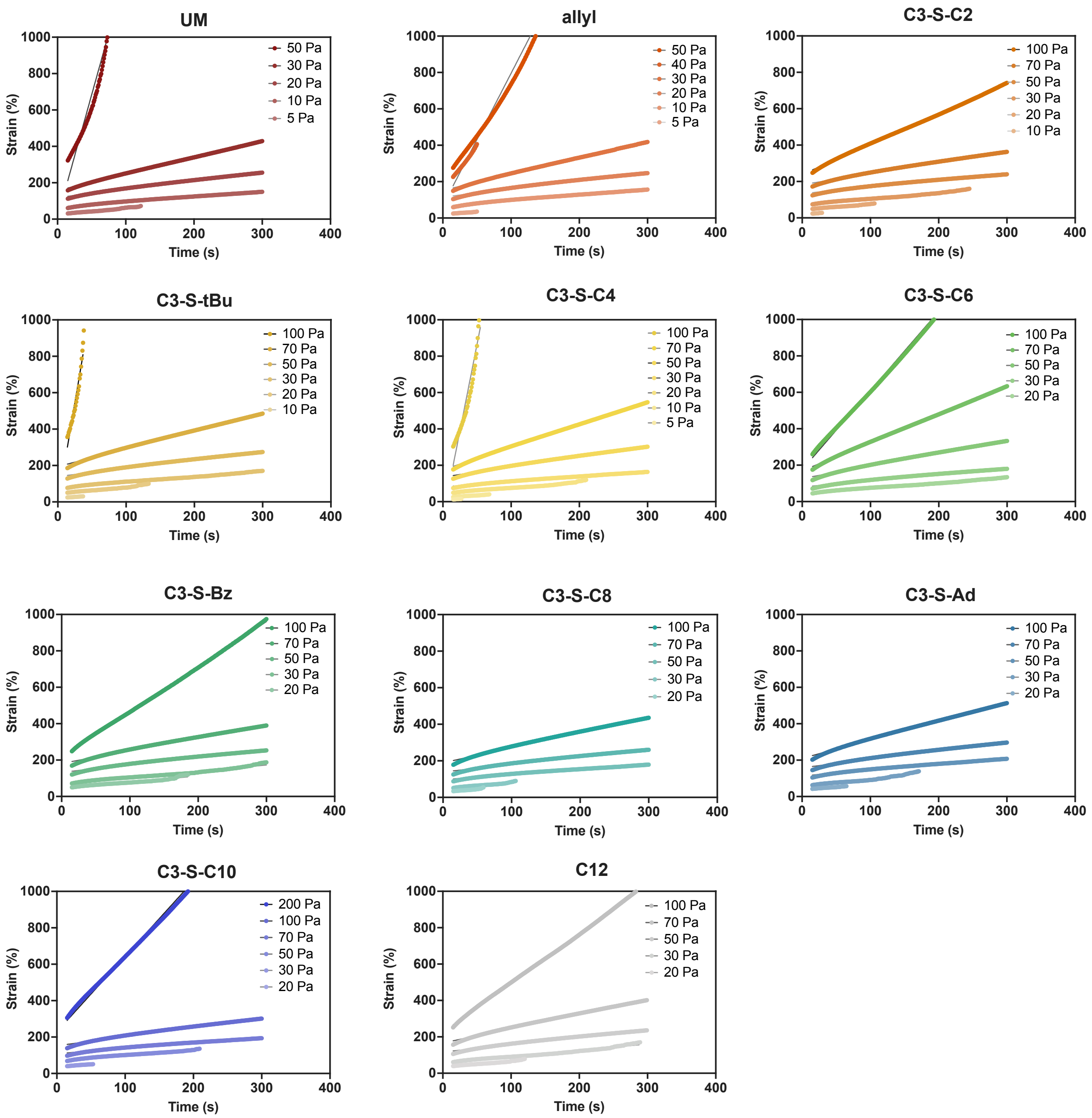

**Figure S 23.** Creep response of for each PNP hydrogel derivative under a range of applied stresses.

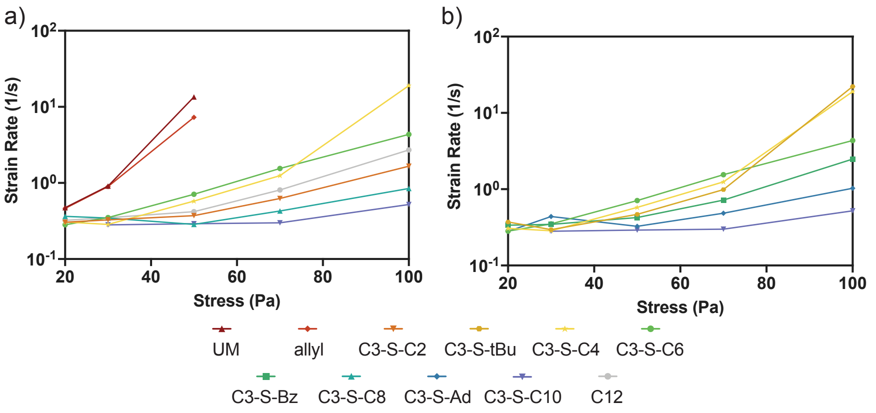

**Figure S 24.** Strain rates at varied stress comparing a) increasing pendant chain length and b) steric effects obtained by plotting slopes of each stress in **Figure S22**.

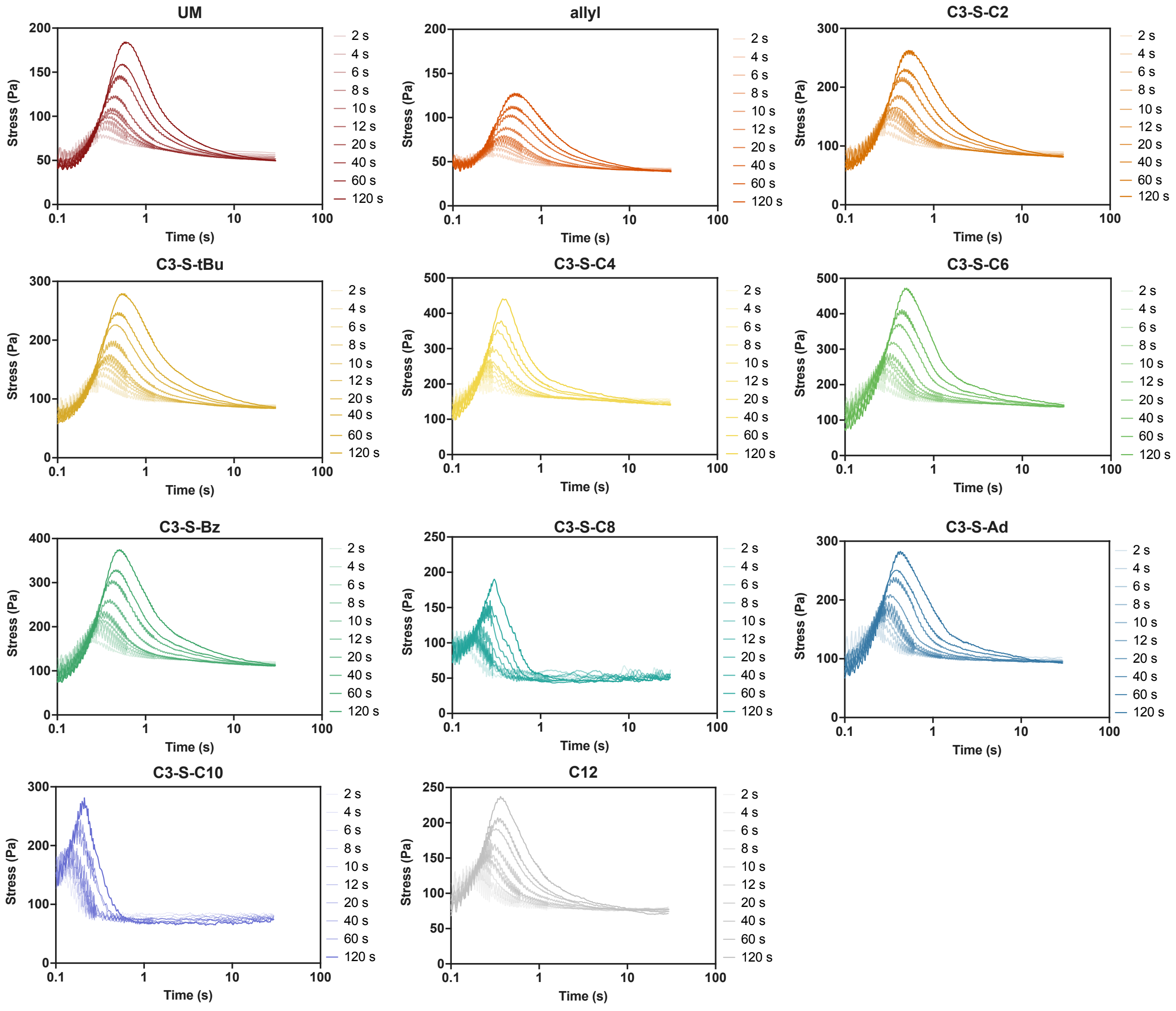

**Figure S 25.** Stress overshoot plots for each PNP hydrogel derivative.

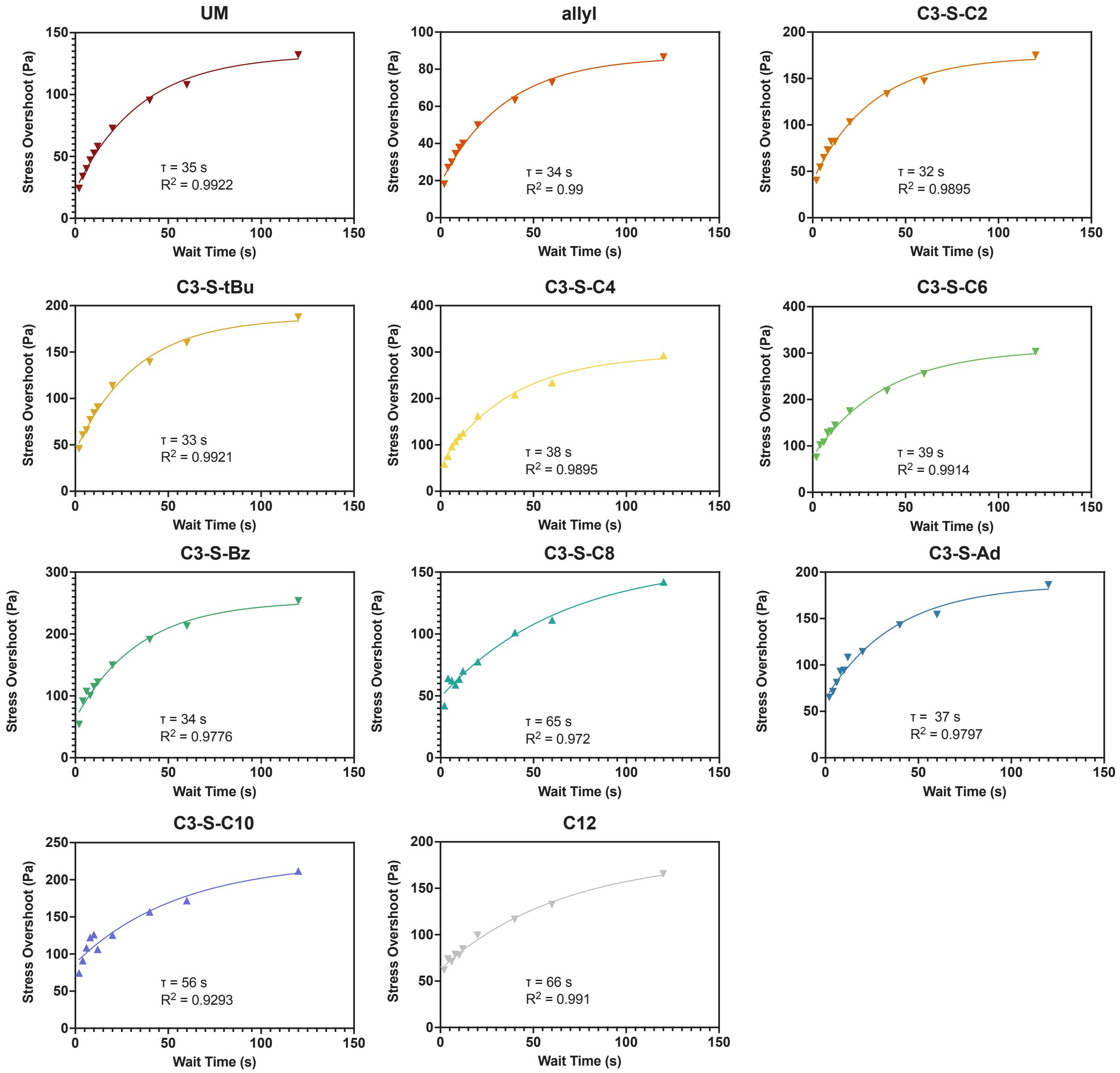

**Figure S 26.** Fitting of stress overshoot plots for each PNP hydrogel derivative to exponential plateaus to obtain characteristic recovery times plotted in **Figure 4d-e**.

### 4. Excision of PNP Depots

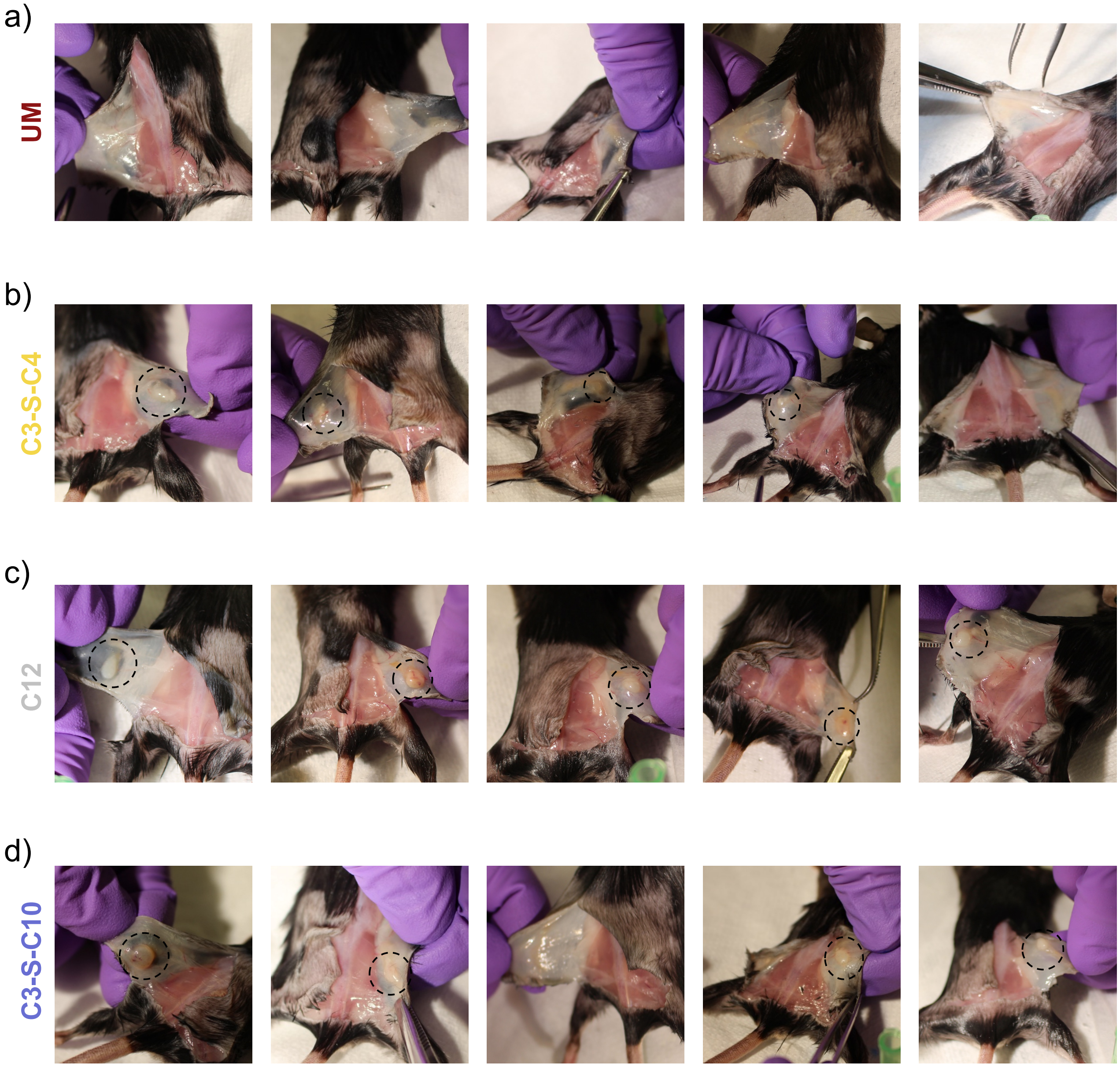

**Figure S 27.** Photographs taken during hydrogel excisions 70 days post-injection. Remaining depots are outlined with black circles.
